# Islet-intrinsic sex differences in inflammatory signaling contribute to autoimmune diabetes susceptibility

**DOI:** 10.1101/2025.10.06.680429

**Authors:** Kierstin L. Webster, Armando A. Puente, Jacob R. Enriquez, Soumyadeep Sarkar, Ernesto S. Nakayasu, Bobbie-Jo M. Webb-Robertson, Lisa M. Bramer, Kayla Figatner, Sarida Pratuangtham, Wenting Wu, Carmella Evans-Molina, Hubert M. Tse, Saptarshi Roy, Jon D. Piganelli, Stephen R. Hammes, Sarah A. Tersey, Raghavendra G. Mirmira

**Affiliations:** Department of Medicine and the Diabetes Research and Training Center, The University of Chicago, Chicago, IL, USA; Biological Sciences Division, Pacific Northwest National Laboratory, Richland, WA, USA; Center for Diabetes and Metabolic Diseases, Indiana University School of Medicine, Indianapolis, IN, USA; Roudebush Veterans Affairs Medical Center, Indianapolis, IN, USA; Department of Microbiology, Molecular Genetics, and Immunology, University of Kansas Medical Center, Kansas City, KS, USA; Division of Endocrinology, Department of Medicine, University of Rochester School of Medicine and Dentistry, Rochester, NY, USA

**Keywords:** Sex differences, estradiol, type 1 diabetes, islet, inflammation, autoimmunity

## Abstract

Whereas most autoimmune diseases exhibit female predominance, type 1 diabetes (T1D) occurs more frequently in males after puberty, suggesting a role for sex hormones in disease modification. Because islet β cells actively shape local immune responses, we hypothesized that sex-specific islet responses to inflammation contribute to this disparity. Using transcriptomic and proteomic analyses of human islets from male and female donors, we found that male islets exhibit a more aggressive response to proinflammatory cytokines, characterized by greater induction of interferon signaling and suppression of developmental signaling compared to female islets. Treatment of human islets and mouse β cells with the sex hormone 17β-estradiol (E2) suppressed inflammatory signaling and markers of β-cell maturity while enhancing developmental gene programs. Complementary studies in non-obese diabetic (NOD) mice showed that E2 treatment reduces diabetes incidence and limits progression to severe insulitis. Islet single-cell RNA sequencing revealed that E2 treatment of NOD mice suppresses interferon signaling, chemokine production, and antigen presentation in β cells, while reducing activation and cytotoxicity pathways in immune cells. In co-culture studies in vitro, E2 pretreatment of mouse islets reduces subsequent activation of T cells, and in an aggressive adoptive transfer model in vivo, E2 pretreatment of the recipient mice was found to attenuate hyperglycemia. These findings support a model in which E2-mediated β-cell reprogramming reduces β-cell immunogenicity and promotes local immune tolerance, offering mechanistic insight into sex-biased T1D susceptibility.

## INTRODUCTION

A remarkable 80% of Americans living with autoimmune diseases are female and many autoimmune diseases (e.g., Sjögrens, lupus, multiple sclerosis) show striking female predominance (1–3). Incomplete X chromosome inactivation of immune-related genes and the proinflammatory effects of estradiol are thought to enhance immune responses in females, conferring protection against infection and cancer, but increasing susceptibility to autoimmunity (4–6). However, the distribution of type 1 diabetes (T1D), a disorder caused by autoimmune destruction of insulin-producing β cells, challenges these prevailing notions. While T1D is diagnosed in roughly equal proportions by sex in children, its incidence is about 1.5 times higher in males than in females after puberty (7–9). The mechanistic underpinnings of the male sex bias in T1D have been largely unexplored.

It is now well established that T1D arises not solely from immune dysregulation but from a bidirectional interaction between immune cells and islet β cells, in which β-cell-intrinsic stress responses contribute actively to establishing a pro-autoimmune islet microenvironment (10).

This process is further shaped by underlying genetic susceptibility (including variants in HLA, preproinsulin, and other risk loci) and by environmental factors that precipitate disease onset (11, 12). Viral infection, which can trigger cellular interferon responses, has been suggested as a potential initiating environmental insult (13, 14). A transcriptional interferon signature has been observed in the blood and pancreas of islet autoantibody-positive individuals at risk for T1D (15, 16). Type 1 and 2 interferons (IFN-α and IFN-γ, respectively) not only activate pathogenic features in immune cells but also augment immunogenic features of β cells. Endoplasmic reticulum stress, formation of neoantigens, and the hyperexpression of major histocompatibility I (MHC I) molecules, among others, are known interferon responses that have been observed in β cells in T1D (17–19). In this regard, recent studies in mouse models suggest that attenuating inflammatory signaling pathways in β cells can curb the subsequent activation of adaptive immunity and thereby protect against diabetes development (20, 21). Thus, measures that support the health and resiliency of β cells in the face of inflammation represent a complementary approach to immune-targeting therapies to delay or prevent T1D.

Several lines of evidence suggest that female sex is protective of β-cell health and function. Clinically, females show a lower incidence of T1D and type 2 diabetes (T2D), both of which involve the failure and/or death of β cells (8, 22). Among individuals diagnosed with T1D, females exhibit higher residual C-peptide levels, an indicator of higher residual β-cell function (23, 24). Single-cell RNA sequencing of human islets from nondiabetic donors reveals that even at baseline, male β cells exhibit amplified signatures of ER and oxidative stress compared to female β cells (25). Whether sex hormones or other characteristics mediate these relative female protections is not entirely clear. It has long been observed, however, that the sex hormone 17β-estradiol (E2) has insulinotropic effects on the β cell (26). Mounting evidence suggests that E2 protects β cells from proapoptotic injury following inflammatory, oxidative, and glucolipotoxic stressors (27–29). We hypothesized that the relative protection of post-pubertal females from T1D reflects sex-based differences in islet responses to inflammatory stress. To test this hypothesis, we employed bulk RNA sequencing and proteomics to compare cytokine-induced responses in human islets from male and female donors. Additionally, we utilized the NOD mouse model to investigate the influence of sex hormones on islet inflammation and diabetes development.

## RESULTS

### Transcriptomic response to inflammation in human islets differs by sex in a discovery dataset

To investigate if the qualitative or quantitative scope of the islet response to inflammation is shaped by sex, we first reanalyzed a publicly available bulk RNA sequencing dataset from human islets treated with proinflammatory cytokines (PIC) IL-1β and IFN-γ (30). The combination of cytokines IL-1β and IFN-γ has been shown to mimic the microenvironmental exposure sustained by islets in early T1D, where the combined effects of innate and adaptive immune cell infiltration lead to the release of these inflammatory mediators (31, 32). These human islets were collected from 10 non-diabetic cadaveric donors (N=6 males and 4 females); a subset of islets from each donor was left untreated as a control (vehicle) and a subset was treated with PIC for 24 hours before RNA isolation and sequencing (**Figure 1A**). Notably, this publicly available dataset was used as a “discovery” approach, recognizing that the donor clinical characteristics were not matched by sex (**Table 1 and Supplemental Table S1**).

**Figure 1:**
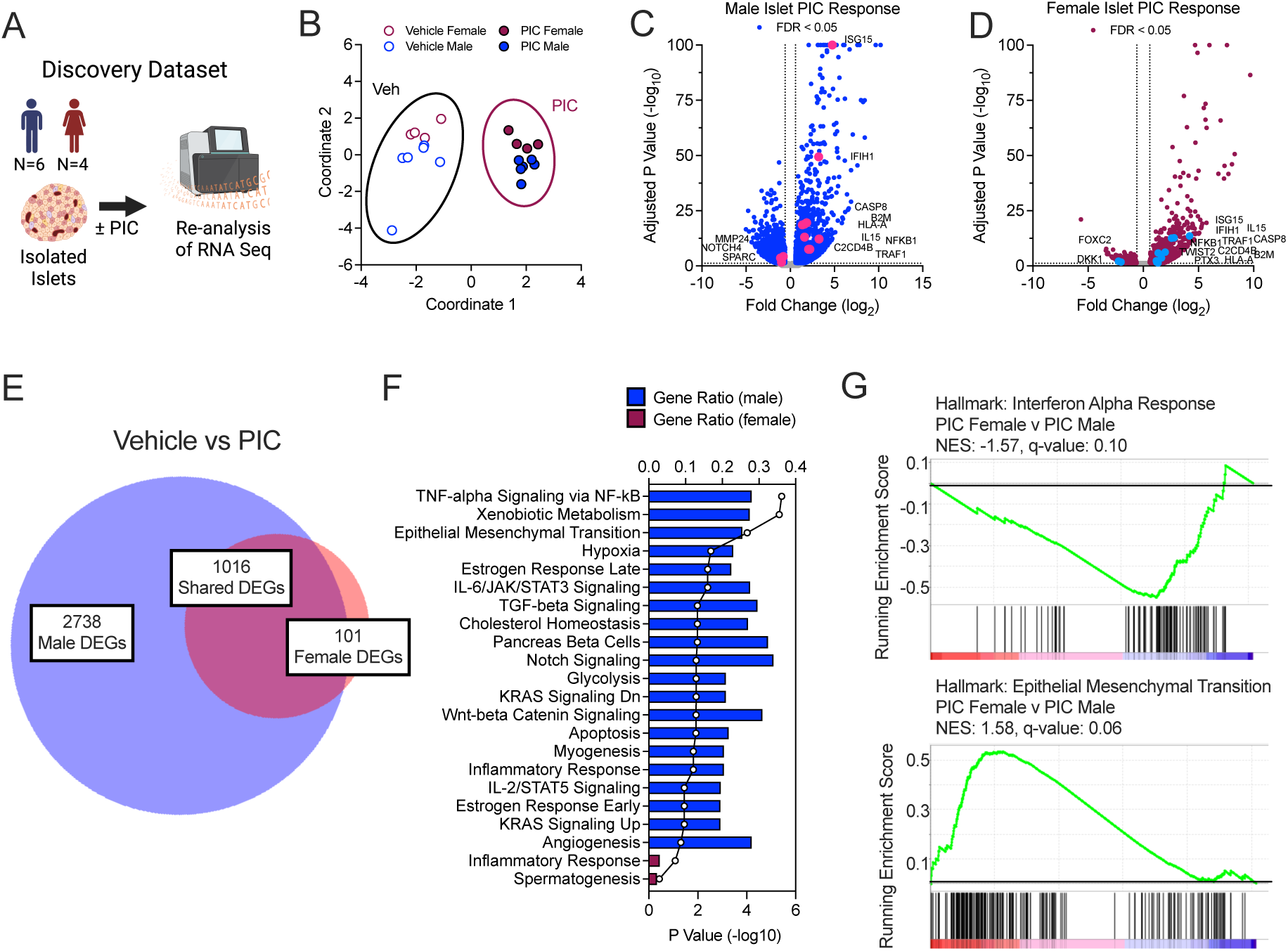
A discovery dataset revealed sexually dimorphic human islet responses to proinflammatory cytokines. Human islets were treated with proinflammatory cytokines (PIC: IL1-β and IFN-γ) and subjected to RNA-sequencing. (**A**) Experimental design. (**B**) Principal component analysis plot. (**C**) Volcano plot of differentially expressed genes in male islets treated with PIC compared to vehicle. Labeled genes highlighted in *red*. (**D**) Volcano plot of differentially expressed genes in female islets treated with PIC compared to vehicle. Labeled genes highlighted in *blue*. (**E**) Shared and unique differentially expressed genes identified in male and female islets treated with and without PIC. (**F**) Hallmark pathway analysis of sex-specific DEGs. (**G**) GSEA of Hallmarks: Interferon Alpha Response (*top panel*) and Hallmarks: Epithelial Mesenchymal Transition (*bottom panel*) in islets from male donors treated with PIC compared to islets from female donors treated with PIC. Islets from human donors: N=6 males and N=4 females.

**Table 1:** Average donor characteristics for discovery and validation human islet samples.

|  | Discovery |  |  | Validation |  |  |
| --- | --- | --- | --- | --- | --- | --- |
|  | Male | Female | P value | Male | Female | P value |
| **Sex (n value)** | 6 | 4 | -- | 10 | 10 | -- |
| **Age (years)** | 26.5 ± 2.8 | 43.0 ± 3.9 | **0.0075** | 43.3 ± 5.0 | 46.4 ± 3.8 | 0.6298 |
| **BMI (kg/m<sup>2</sup>)** | 27.5 ± 2.0 | 31.0 ± 2.1 | 0.2615 | 29.1 ± 1.5 | 27.7 ± 1.8 | 0.5865 |
| **HbA1c (%)** | 5.2 ± 0.1 | 5.6 ± 0.1 | **0.0153** | 5.3 ± 0.1 | 5.3 ± 0.1 | 0.8874 |

Including islets from all 10 donors, principal component analysis revealed that gene expression changes largely stratified by treatment with PIC (**Figure 1B**); upon filtering for genes with a fold-change ≥1.5 or ≤-1.5 and an adjusted p-value <0.05, 3793 differentially expressed genes (DEGs) were identified in response to PIC (**Supplemental Figure S1A**). By overrepresentation analysis, these DEGs mapped to expected inflammatory Human Molecular Signatures Database (MsigDB) Hallmarks terms, including IFN-γ and IFN-α responses, Allograft Rejection, JAK/STAT signaling, and Apoptosis, among others (**Supplemental Figure S1B**). Strikingly, separating the analysis by sex revealed that 3754 genes were differentially expressed in PIC-treated male islets compared to vehicle, whereas only 1117 were differentially expressed in

PIC-treated female islets (compared to vehicle) (see volcano plots in **Figure 1C and D**)—a nearly 3.5-fold difference between sexes. A notable 2738 DEGs were unique to male islets, whereas only 101 were unique to female islets (**Figure 1E** and all identified genes in **Supplemental Table 2**). Functional pathways represented among the 2738 male-specific DEGs related to inflammation and cell death (TNF-α Signaling via NF-kB, IL-6/JAK/STAT3 Signaling, Apoptosis) and cell identity and development (Notch Signaling, Wnt-β catenin Signaling, Pancreas Beta Cells, Epithelial-Mesenchymal Transition) (**Figure 1F**). Among the 101 female-specific DEGs were a small number of genes related to Inflammatory Response (**Figure 1F**).

Many of the DEGS unique to male islets were aligned with the interferon response (see volcano plot, **Figure 1C**) and in those unique to female islets were aligned with the epithelial-to-mesenchymal transition (EMT) and extracellular matrix (see volcano plot, **Figure 1D**). To assess how gene sets associated with these processes might be enriched in each sex (regardless of statistical changes to specific genes), we next compared male and female islet transcriptomes directly using gene set enrichment analysis (GSEA). PIC-treated male islets showed enrichment of the IFN-α response compared to PIC-treated female islets (**Figure 1G**). IFN-α is known as a ‘key trigger’ for T1D (33), and several genes involved in IFN-α response are considered T1D candidate genes, including *IFIH1, PTPN22,* and *TYK2* (34, 35). Notably, the “leading-edge” genes—those contributing most strongly to the enrichment of this pathway in male islets-included T1D candidate gene *IFIH1*, as well as *ISG15, IL15,* and *CASP8,* each implicated in T1D pathogenesis (**Supplemental Figure S1C**) (36–40). Female islets, on the other hand, showed an enrichment in genes involved in the EMT pathway compared to male islets (**Figure 1G, Supplemental Figure S1D**). EMT has been observed in β cells undergoing dedifferentiation in prolonged cell culture and also in models of diabetes (41–44). While dedifferentiation is thought to be a mechanism of β cell failure in T2D (45, 46), it has been shown to *protect* β cells from autoimmune destruction in the NOD model of T1D by decreasing expression of autoantigens and MHC I (47, 48).

### A validation dataset recapitulates the sexually dimorphic islet transcriptomic responses to inflammation

The observations that male islets show relative enrichment of inflammatory response pathways and female islets show enrichment of EMT genes in the discovery dataset led us to validate these findings with a prospective collection of human donor islets. We performed bulk RNA sequencing on a cohort of human islets (N=10 males and 10 females) that were well-matched for age, BMI, and HbA1c levels (**Table 1 and Supplemental Table S1**). A subset of islets from each donor was untreated as a control (vehicle), and another was treated with IL-1β and IFN-γ for 18 hours (PIC) (**Figure 2A**), similar to the discovery dataset. When considering samples collectively from all 20 donors, principal component analysis indicated that gene expression changes segregated primarily by response to PIC (**Figure 2B**). 2134 DEGs were identified in response to PIC (FC >1.5, adj-p-value<0.05) (volcano plot, **Supplemental Figure S2A**). Overrepresentation analysis showed that these DEGs belong to many of the same MSigDB Hallmarks terms noted in the discovery dataset, including IFN-α and IFN-γ Response, Allograft Rejection, IL-2/STAT5 Signaling, Apoptosis, and Epithelial Mesenchymal Transition, among others (**Supplemental Figure S2B**). When considering samples by sex, the number of DEGs in PIC-treated male islets (compared to vehicle) was again larger than that in PIC-treated female islets (2386 in males; 1765 in females) (see volcano plots in **Figure 2C and D** and all identified genes in **Supplemental Table 3**), and males showed over 3.5 times more sex-specific DEGs than females (864 in males; 243 in females) (**Figure 2E**). Male-specific DEGs related to pro-inflammatory signaling (i.e. TNF-α Signaling via NFKB), KRAS Signaling, Apical Junction, and Angiogenesis, as well as EMT (**Supplemental Figure S2C**). Female-specific DEGs, on the other hand, related to Estrogen Responses and EMT (**Supplemental Figure S2C**).

**Figure 2:**
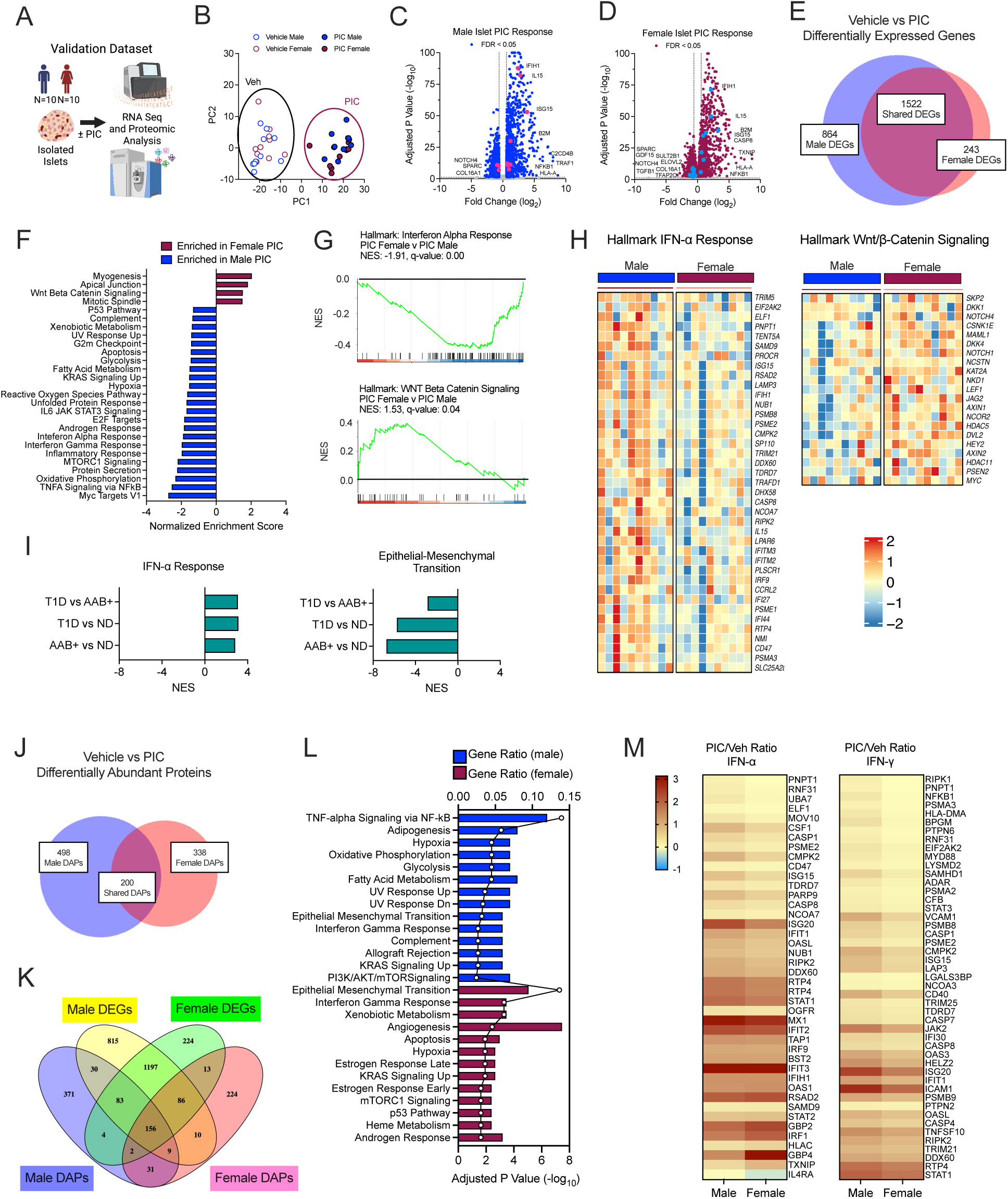
A validation dataset confirmed sexually dimorphic human islet responses to proinflammatory cytokines. Human islets were treated with proinflammatory cytokines (PIC: IL1-β and IFN-γ) and subjected to RNA-sequencing and proteomics. (**A**) Experimental design. (**B**) Principal component analysis plot. (**C**) Volcano plot of differentially expressed genes in male islets treated with PIC compared to vehicle. Labeled genes highlighted in *red*. (**D**) Volcano plot of differentially expressed genes in female islets treated with PIC compared to vehicle. Labeled genes highlighted in *blue*. (**E**) Shared and unique differentially expressed genes identified in male and female islets treated with and without PIC. (**F**) GSEA comparing PIC-treated male islets and PIC-treated female islets. (**G**) GSEA of Hallmarks: Interferon Alpha Response (top panel) and Hallmarks: WNT Beta Catenin Signaling (*bottom panel*). (**H**) Heatmaps depicting expression of leading edge genes in PIC-treated male and female islets for Interferon Alpha Response GSEA (left panel) and WNT Beta Catenin Signaling GSEA (*right panel*). (**I**) GSEA of Hallmarks: Interferon Alpha Response (*left panel*) and Hallmarks: Epithelial-Mesenchymal Transition (*right panel)* in β cells from Control, AAB+, and T1D cadaveric donors from the Human Pancreas Analysis Program’s PancDB database. Control: non-diabetic donors, N=15; AAB+: single and double autoantibody positive donors, N=10; T1D: donors with T1D, N=9. (**J**) Shared and unique differentially abundant proteins (DAPs) identified in male and female islets from the validation cohort treated with and without PIC. (**K**) Venn diagram depicting overlap between differentially expressed genes and differentially abundant proteins identified in response to PIC in islets from male and female donors. (**L**) GO: Biological Pathways analysis of sex-specific differentially abundant proteins in response to PIC. (**M**) Heatmaps of fold-change (log2) in protein abundance of proteins in the Interferon Alpha Response (*left panel*) and Interferon Gamma Response pathways (*right panel*) in response to PIC. Islets from human donors: N=10 males and N=10 females.

Using GSEA to compare transcriptomes of PIC-treated male and female islets, we identified several signaling pathways with sex-specific enrichment (**Figure 2F**). Notably, PIC-treated male islets were enriched for IFN-α Response relative to PIC-treated females as observed in the discovery set and also for several related pathways including IFN-γ Response, Unfolded Protein Response, Reactive Oxygen Species, and Apoptosis (**Figure 2F**). The leading-edge genes driving the male enrichment of IFN-α Response genes overlaps largely with those in the discovery set and includes genes implicated in T1D pathophysiology (e.g. *IFIH1, ISG15, IL15,* and *CASP8*) (**Figure 2G**). Female islets were enriched for developmental pathways including Myogenesis, Wnt Beta Catenin signaling, and Apical Junction, each important to the development and maintenance of endocrine cell identities in the islet (49–51) (**Figure 2F**). Genes in these pathways are shared with those in the EMT pathway as observed in the discovery dataset, including NOTCH receptors (*NOTCH2, NOTCH4)*, collagens (*COL6A2, COL16A1*), and other secreted factors involved in tissue remodeling (*TGFB1, BMP1, MMP2)*(**Figure 2G**). Underscoring the relevance of these sexually dimorphic pathways to T1D pathogenesis, GSEA of single cell RNA sequencing on human islet β cells from the Human Pancreas Analysis Program (HPAP) including cells from nondiabetic, autoantibody positive (AAB+), and T1D donors (52–54) revealed that IFN-α Response is enriched in β cells with T1D progression while EMT is suppressed (**Figure 2I**).

### Islet proteomics confirms sexually dimorphic alterations in protein levels with inflammation

To assess how inflammation alters islet protein levels in a sex-specific manner, we performed mass spectrometry-based proteomics on islets from the validation cohort. Proteomics was performed after the same 18 h PIC treatment period on islet extracts from the same samples used for transcriptomics. Male islets showed 698 distinctly altered proteins in the presence of PIC, whereas female islets showed 538 altered proteins (adj-p-value<0.05; **Figure 2J**, and **Supplemental Table 4** for all identified proteins). Male islets showed about 1.5 times more unique differentially abundant proteins in response to PIC than female islets (adj-p-value<0.05; 498 unique to males, 338 unique to females) (**Figure 2J** and **Supplemental Table 4).** Several similarities with the transcriptomics emerge from the proteomics: (a) males showed a greater number of differentially abundant proteins in response to PIC compared to females (**Figure 2J**), (b) there was substantial overlap in PIC responsive genes and proteins (278 genes and proteins overlapped in males, 257 overlapped in females) (**Figure 2K**), and (c) based on overrepresentation analysis, male-specific changes in proteins included the Hallmarks terms TNF-α Signaling via NFKB, KRAS Signaling Up, and EMT, while female-specific changes included the Hallmarks terms Estrogen Response Early, Estrogen Response Late, and EMT, as was shown in transcriptomics (**Figure 2L**). Proteins related to interferon responses, shown to be enriched in male islets at the transcriptomic level, also showed more robust changes in abundance in male islets in response to PIC (**Figure 2M**). Notably, several genes/proteins showing consistent sexually dimorphic responses to PIC at both the transcriptomic and proteomic levels have demonstrated roles in autoimmunity (*NFKB1*, *TRAF1*) (55, 56), β-cell function (*SPARC*, *C2CD4B*) (57–59), and putatively T1D pathophysiology (e.g. *TXNIP*, *GDF15*) (60, 61) (**Supplemental Figure S2D**). Overall, these data suggest that coordinate changes to both proteins and transcripts in the islet in response to PIC occur in a sexually dimorphic manner that could influence disease susceptibility.

### *17β-Estradiol* treatment suppresses inflammatory signaling while stimulating developmental pathway signaling in the islet in vitro

Male and female incidence of T1D diverges after the age of puberty (7, 8), pointing to sex hormones as logical suspects for protection from or predisposition to disease. Among the female-specific DEGs identified in the validation transcriptomic dataset in response to PIC were genes related to estrogen responses, such as *SULT2B1, TFAP2C, MUC1, and ELOVL2* (**Supplemental Figure S2C** and **Supplemental Table 3**). This observation suggests that at least some of the sex-specific differences observed in response to PIC may relate to signaling by 17β-estradiol (E2) in the islet. To investigate how PIC responses may be influenced by E2 signaling, we first examined expression of the genes encoding estrogen receptors ERα and ERβ *(ESR1* and *ESR2*) as well as plasma membrane estrogen receptor GPER1 in our validation RNA sequencing dataset. PIC treatment had no significant effect on *ESR2* transcript levels in either male or female islets (**Figure 3A**). In islets from both sexes, *GPER1* expression was decreased by PIC (**Figure 3A**). Notably, PIC treatment led to reduced *ESR1* levels in male islets, with no effect on *ESR1* in female islets (**Figure 3A**), suggesting the potential that male islets might become desensitized to the effect of E2 through its major receptor in the presence of cytokines. Similar to human male islets, both *Esr1* and *Esr2* were statistically unchanged in T1D-prone NOD male and female mouse islets (**Figure 3B**). In contrast to these data from humans and NOD mice, PIC treatment of murine insulinoma β cells (MIN6) and *C57BL/6J* mouse islets revealed increased *Esr1* expression (**Figure 3C and 3D**), with no significant changes in *Esr2*. Together, these data suggest that islets may alter their receptiveness to E2 signaling by correspondingly altering a relevant E2 receptor (*ESR1/Esr1*) in response to inflammation.

**Figure 3:**
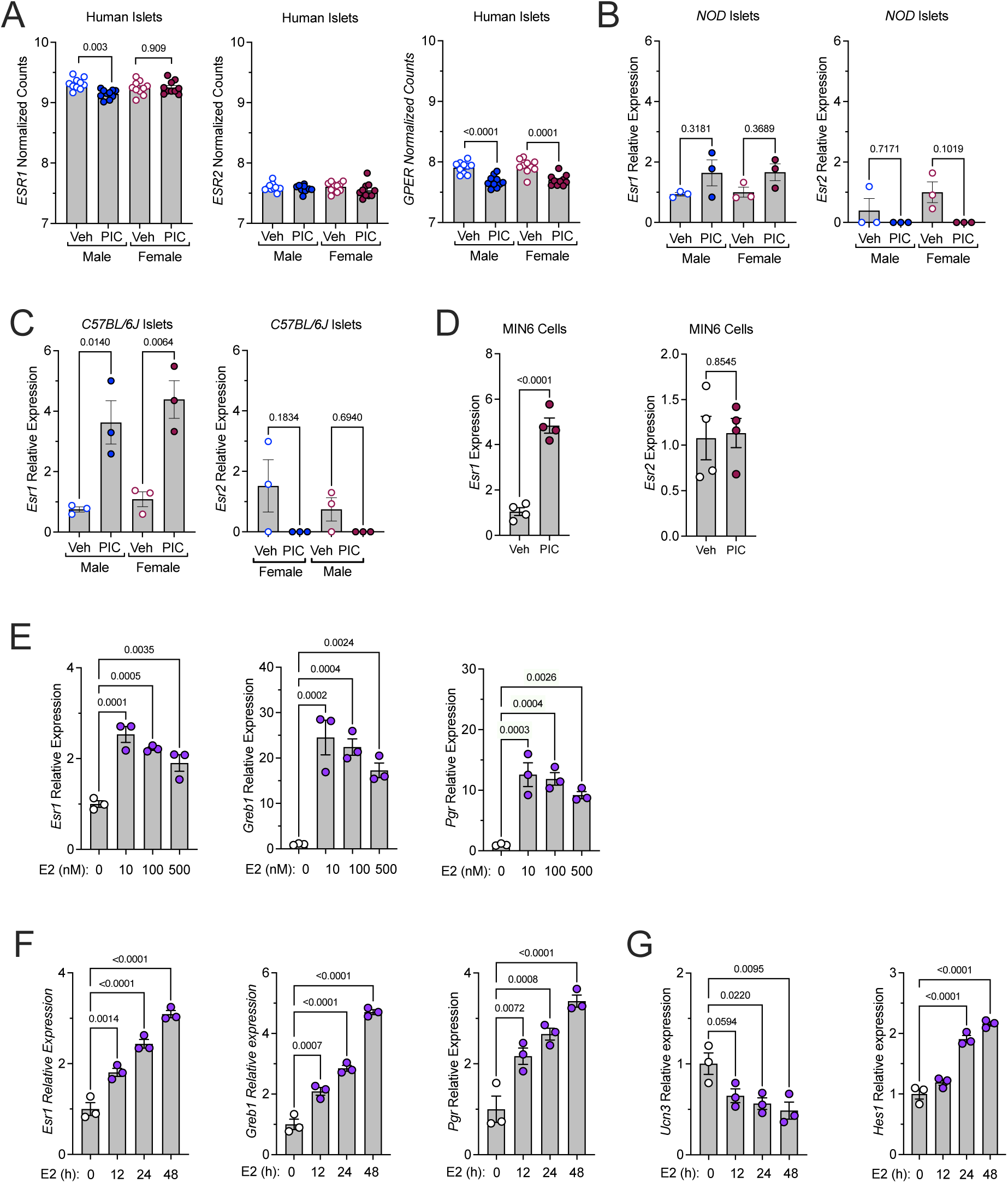
Proinflammatory cytokines alter estrogen receptor β (*ESR1*) but not estrogen receptor β (*ESR2*). Human islets, C57BL/6J islets, NOD islets, and MIN6 β cells were treated with PIC and gene responses were measured. (**A**) *ESR1* (*left panel*), *ESR2* (*middle panel*), and *GPER1* (*right panel*) gene expression values in male and female human islets after treatment with proinflammatory cytokines (PIC: IL1-β and IFN-α). (**B**) *Esr1* (*left panel*) and *Esr2* (*right panel*) gene expression values in male and female C57BL/6J mouse islets after treatment with proinflammatory cytokines for 18h (PIC: IL1-β, IFN-γ, TNF-α). (**C**) *Esr1* (*left panel*) and *Esr2* (*right panel*) gene expression values in male and female NOD mouse islets after treatment with proinflammatory cytokines for 18h (PIC: IL1-β, IFN-γ, TNF-α). (**D**) *Esr1* (*left panel*) and *Esr2* (*right panel*) gene expression values in MIN6 β cells after treatment with proinflammatory cytokines (PIC: IL1-β, IFN-γ, TNF-α). (**E**) *Esr1* (*left panel*), *Pgr* (*middle panel*), and *Greb1* (*right panel*) gene expression values in MIN6 β cells after treatment with different doses of E2. (**F**) *Esr1* (*left panel*), *Pgr* (*middle panel*), and *Greb1* (*right panel*) gene expression values in MIN6 β cells after treatment with a time course of 10 μM E2. (**G**) *Ucn3* (*left panel*) and *Hes1* (*right panel*) gene expression values in MIN6 β cells after treatment with a time course of 10 μM E2.

To assess β-cell responsiveness to E2 in vitro, we first examined murine-derived β cells to identify an appropriate E2 concentration. MIN6 β cells were treated with E2 concentrations ranging from 10 nM to 500 nM, and expression of selected ERα target genes was measured: *Esr1* itself, *Greb1* (encoding an ERα cofactor), and *Pgr* (encoding progesterone receptor).

Expression of all 3 genes was significantly increased at all concentrations tested after 18 hours (**Figure 3E**). Treatment for up to 48 hours with the lowest concentration (10 nM E2) produced more pronounced increases in ERα response gene expression (**Figure 3F**). Because we observed in our discovery and validation datasets that female islets exhibit enriched signaling in developmental pathways, we hypothesized that E2 exposure might alter the state of β-cell maturity. Over the 48-hour time course, expression of β-cell maturity gene *Ucn3* declined while expression of *Hes1*, an effector of Notch signaling associated with β-cell dedifferentiation (62), increased (**Figure 3G**). Overall, these data suggest that β cells respond to E2 treatment, and that E2 may decrease β-cell maturity while activating developmental pathways observed to be enriched in female human islets relative to male islets.

To investigate the effect of E2 signaling on human islets in vitro, a subset of islets from each donor from the validation cohort (**Table 1**) was treated for 18 hours with 10 nM E2, after which both RNA and protein were collected (**Figure 4A)**. The RNA from the E2-treated samples was bulk sequenced alongside the vehicle and PIC treated samples discussed earlier. The effects of this acute E2 treatment were much more subtle than those of acute PIC; thus, for this analysis the looser filtering criteria of fold-change ≥1.25 or ≤-1.25 and p-value < 0.05 were used. By sex, the transcriptomic responses to E2 were starkly different. While males showed 235 DEGs in response to E2, females showed 171 DEGs, with only 17 genes overlapping between sexes (**Figure 4B)**. Among the overlapping response genes was canonical ERα target *GREB1* (**Figure 4B**). While the individual DEGs were highly disparate between sexes, overrepresentation analysis and GSEA-strategies, which focus on coordinated changes within a functional pathway rather than specific genes, highlighted some functional overlap. By overrepresentation analysis, pathways related to E2 responses (Estrogen Response Early and Late) and inflammatory processes (e.g. IFN-γ Response, Allograft Rejection) were among those modulated in islets of both sexes with E2 treatment (**Figure 4D**). Further, GSEA revealed that E2 treatment suppressed the IFN-γ Response pathway in islets from both sexes (**Figure 4E and 4F**), suggesting that prior exposure to E2 in vivo may indeed help to explain the diminished IFN signatures observed in female islets compared to male islets (**Figures 1G and 2G**).

**Figure 4:**
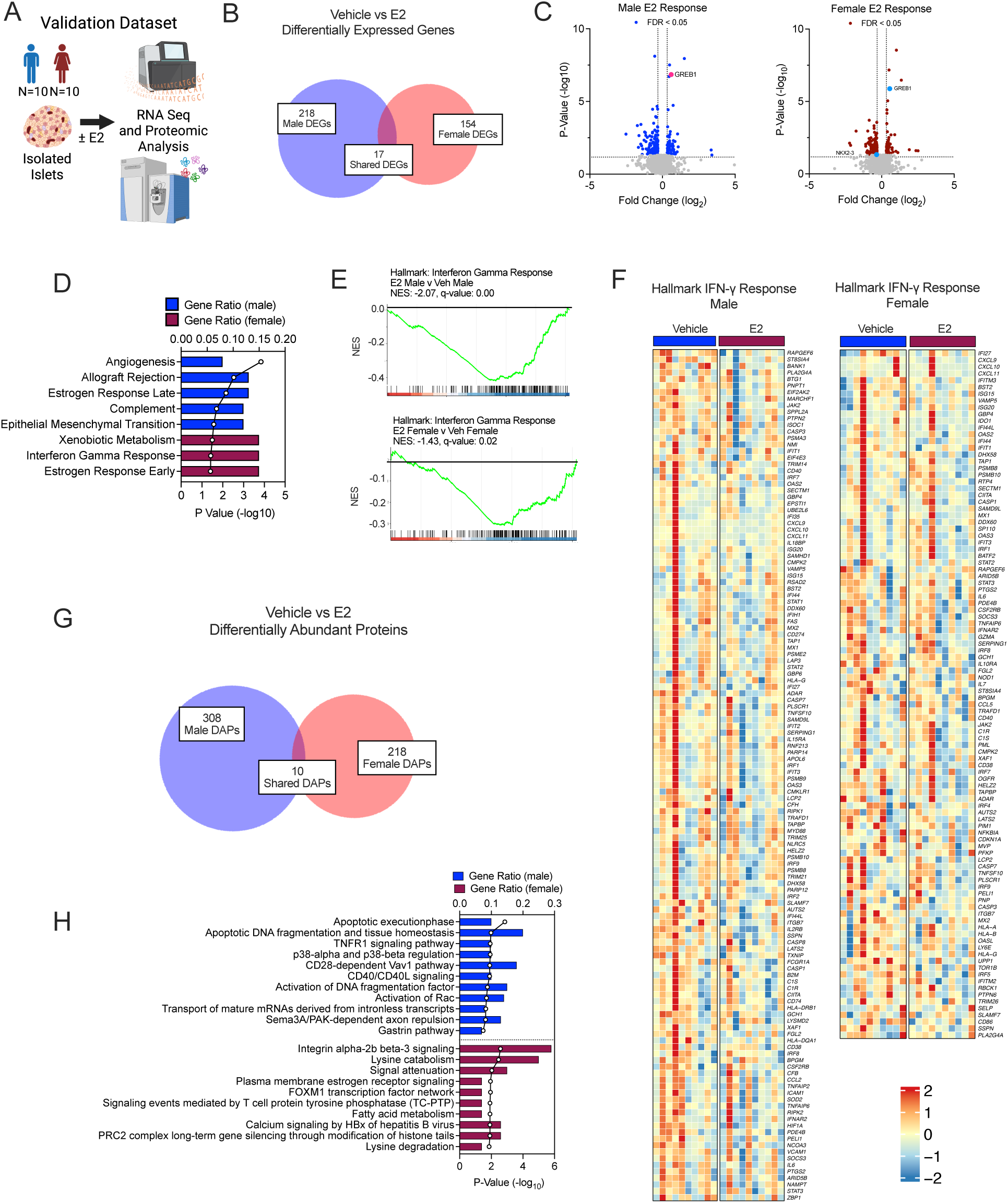
RNA sequencing and proteomics revealed sexual dimorphic human islet responses to estradiol. Human islets were treated with vehicle or estradiol (E2) for 18h and subjected to RNA-seq and proteomics. (**A**) Experimental design. (**B**) Shared and unique differentially expressed genes (DEGs) identified between male and female islets treated with and without PIC. (**C**) Volcano plots of differentially expressed genes in male islets (*left panel*) and female islets (*right panel*) treated with PIC compared to vehicle. (**D**) Hallmark pathway analysis of sex-specific differentially expressed genes. (**E**) GSEA of Hallmark: Interferon Gamma Response (*bottom panel*) in islets from male donors (top panel) and female donors (*bottom panel*) treated with E2 compared to vehicle. (**F**) Heatmaps depicting expression of the leading edge genes from Interferon Gamma Response GSEA in Vehicle vs. E2 conditions in islets from male donors (left panel) and female donors (*right panel*). (**G**) Shared and unique differentially abundant proteins (DAPs) identified in male and female islets treated with and without E2. (**H**) Bioplanet analysis of sex-specific differentially abundant proteins. Islets from human donors: N=10 males and N=10 females.

Mass spectrometry-based proteomics showed that the E2 response was also highly sexually dimorphic at the protein level, with male islets showing 318 differentially abundant proteins and female islets showing 228, with only 10 shared proteins between the sexes (**Figure 4G and Supplemental Table 4**). Male islets showed changes in apoptotic signaling pathways (e.g. Apoptotic Execution Phase) in response to E2, a signal that was not detectable in the female response (**Figure 4H**). Instead, female changes included several metabolic pathways and, of note, Plasma Membrane Estrogen Receptor Signaling (**Figure 4H**). In summary, the estrogen-stimulated β-cell and islet data in vitro suggest that E2 incites potentially protective signaling in both male and female islets, such as decreasing IFN-γ signaling. The overall lack of overlap between differentially expressed genes and proteins in response to E2 by sex in human islets might reflect different modalities of E2 signaling, perhaps driven by E2 sensing by different receptor types (i.e. ERα vs. ERβ isoforms, or nuclear vs. membrane-associated receptor pools), regulation via different estrogen receptor co-regulators, or by differential accessibility of estrogen response elements on the DNA between sexes. These data raise the possibility that E2-stimulated signaling in the islet cells may contribute to female protection from autoimmune destruction.

### Estradiol treatment protects against diabetes development in male and female NOD mice

To investigate the protective effects of E2 on islet function and its impact on autoimmune diabetes development, we next turned to the non-obese diabetic (NOD) mouse model, which develops spontaneous autoimmune diabetes through an interplay between β cells, innate immune cells, and adaptive immune cells (63). Male and female NOD mice were subcutaneously implanted with placebo-or 0.5 mg E2-containing pellets formulated to deliver their contents over 90 days (**Figure 5A**). Treatment was initiated at 6 weeks of age, a time when NOD mice begin to exhibit insulitis but have not yet developed diabetes (64), and diabetes incidence was monitored up to 25 weeks of age. E2-treated male mice were completely protected from diabetes development during the entire study period (**Figure 5B**). Female mice implanted with E2 were completely protected from diabetes onset during the 90 days of pellet activity, while approximately 50% of placebo-treated females developed diabetes during the same period (**Figure 5B**). Twenty-five percent of E2-treated females went on to develop diabetes after the expiration of the E2 pellet.

**Figure 5:**
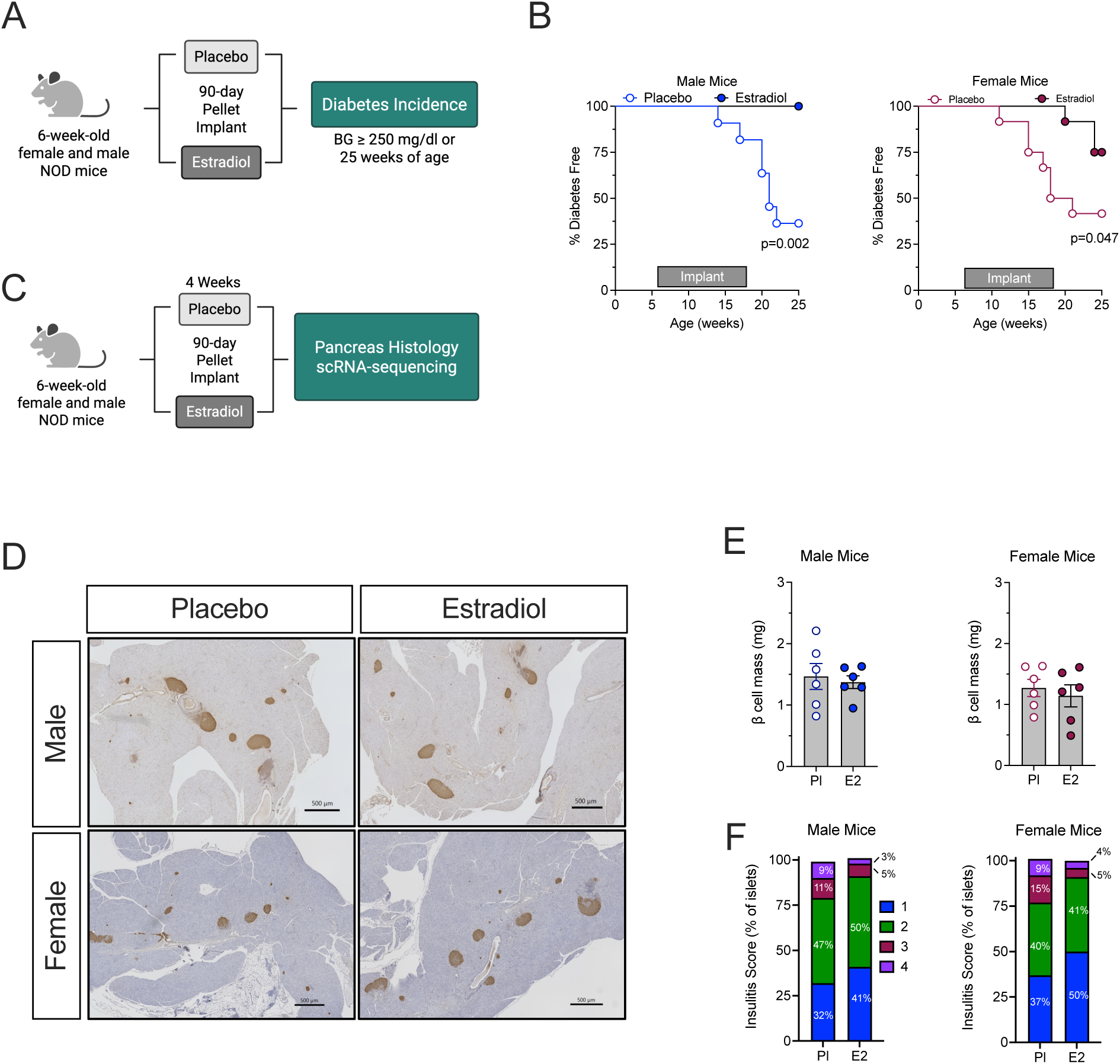
Estradiol treatment protected NOD mice from autoimmune diabetes. Male and female pre-diabetic NOD mice were treated with placebo or E2 90-day pellet implants. (**A**) Experimental design for diabetes incidence studies. (**B**) Diabetes incidence in male (*left panel*) and female (*right panel*) NOD mice treated with placebo and E2. N=10-12 mice per group. (**C**) Experimental design for short-term experiments. (**D**) Representative images of pancreata from placebo and E2-treated NOD male (*top panels*) and female (*bottom panels*) mice stained for insulin (*brown*), and counterstained with hematoxylin (*blue*). Scale bars: 500 μm. (**E**) β-cell mass in E2-treated NOD male (*left panel*) and female (*right panel*) mice. N=6 mice per group. (**F**) Insulitis scores in islets from E2-treated NOD male (*left panel*) and female (*right panel*) mice. N=6 mice per group. Statistical tests: Mantel-Cox: **B**. Unpaired t-test: **E and F**. Data are represented as mean ± SEM.

To explore the effects of E2 on islets during the prediabetic period, pancreas and serum samples were collected from mice treated with either E2 or placebo pellets from 6 to 10 weeks of age (**Figure 5C**). E2 pellet implantation led to an expected increase in circulating E2 levels compared to placebo implantation (**Supplemental Figure 3A**). E2 treatment did not change average β-cell mass after 4 weeks in either male or female mice (**Figure 5E**). Whereas E2 treatment did not reduce overall infiltration of islets by immune cells, it did decrease the average proportion of severely insulitic islets, defined as those in which immune infiltrates cover 50% or greater of islet area (scores 3 and 4) (**Figure 5F**). These findings suggest that, during the early stages of disease, E2 does not prevent immune cell entry into the islet but may instead modulate initial β cell–immune cell interactions in a manner that limits progression to more severe insulitis.

### Estradiol treatment promotes a tolerogenic phenotype in β cells

To assess the quantities and transcriptional profiles of islet endocrine and immune cells after treatment of prediabetic NOD mice with placebo or E2 pellets, we next performed single-cell RNA sequencing (scRNAseq) on islets isolated from 10-week-old female NOD mice after 4 weeks of treatment. Three samples were sequenced per treatment group, each comprising islets pooled from 2 animals, and cell types were identified by the presence of transcripts known to be enriched in each type (**Supplemental Figure S3B**). Islet cell populations identified included all major islet endocrine cell types (β, α, δ, and PP cells), infiltrating immune cells (T and B lymphocytes, myeloid cells), as well as some supporting cell types (acinar, ductal, and mesenchymal cells, erythrocytes) (**Figure 6A**).

**Figure 6:**
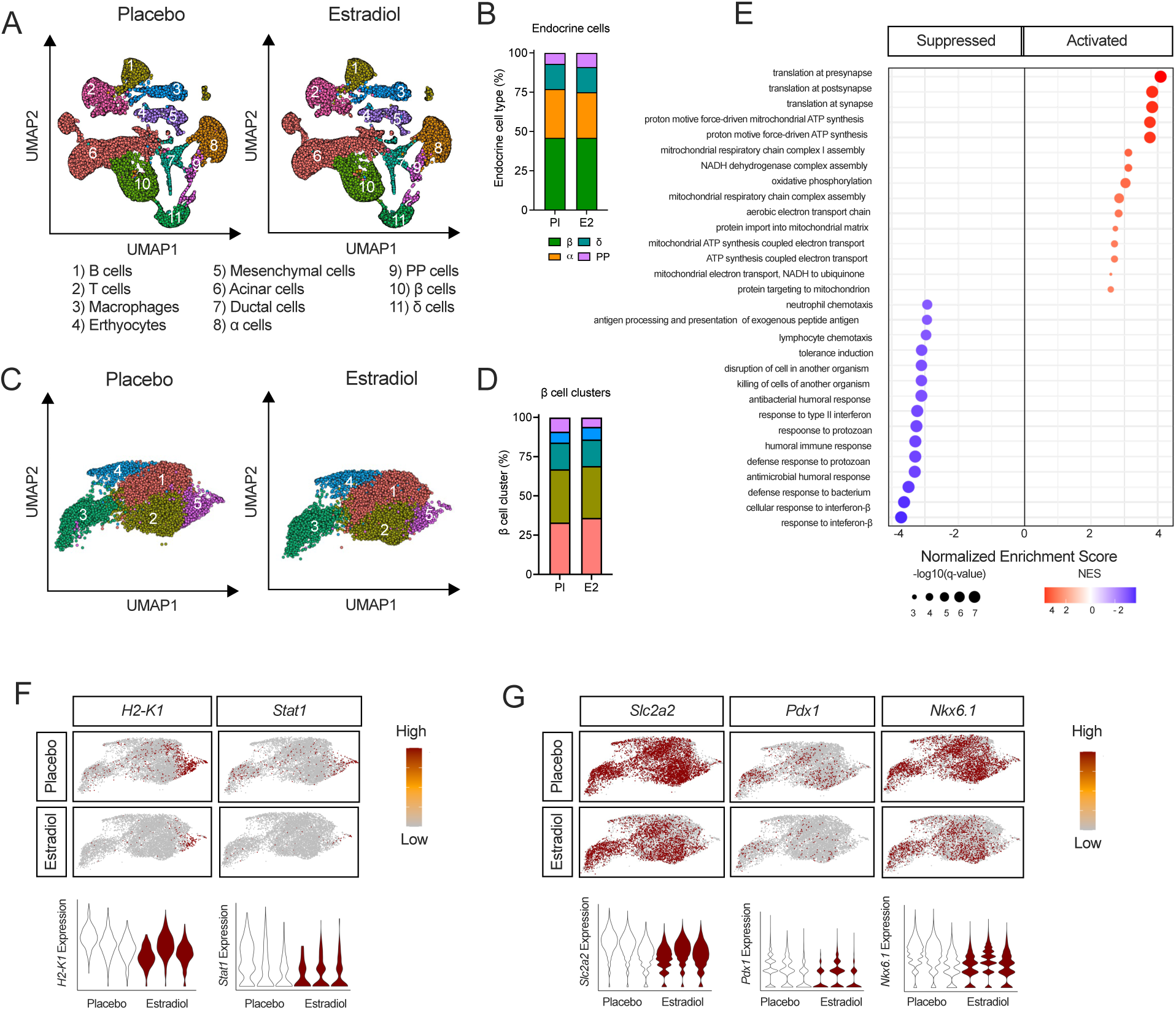
Single-cell RNA-seq revealed β cell differences after estradiol treatment of female NOD mice. Prediabetic female NOD mice were treated with placebo or E2 pellet implants for 4 weeks and isolated islets were subjected to scRNA-seq. N=3 biological replicates per group. (**A**) Uniform manifold approximation and projection (UMAP) embeddings of merged single-cell RNA sequencing profiles from islets isolated from mice treated with placebo (*left panel*) or E2 (*right panel*) colored by identified cell types. (**B**) Percent of α, β, δ, and PP cells identified within the islet endocrine cell clusters. (**C**) UMAP embeddings of the β cell clusters of islets isolated from mice treated with placebo (*left panel*) or E2 (*right panel*). (**D**) Percent of β cells identified within the islet β cell clusters. (**E**) GSEA of the β cell clusters. (**F**) Feature plots of H2-K1 and Stat1 gene expression in β cell population (*top panels*). Quantification of H2-K1 and Stat1 gene expression in β cell cluster #5 (*bottom panels*). (**G**) Feature plots of Slc2a2, Pdx1, and Nkx6.1 gene expression in the β cell population (*top panels*). Quantification of Slc2a2, Pdx1, and Nkx6.1 gene expression in all β cell clusters (*bottom panels*).

Gene set enrichment analysis (GSEA) of the β-cell population revealed that, although β-cell abundance was not altered by E2 treatment (**Figure 6B and 6C**), the transcriptomic landscape was substantially reshaped. E2 suppressed key immune-related pathways, including Type I and Type II Interferon responses, Antigen Processing and Presentation, and Lymphocyte Chemotaxis (**Figure 6E**). By contrast, pathways upregulated by E2 were predominantly associated with protein synthesis and mitochondrial ATP production—processes essential for preserving β cell biosynthetic capacity, including insulin secretion. IFN-stimulated genes expressed at significantly lower levels in β cells from E2-treated mice included the known T1D risk gene *Ifih1*, as well as *Stat1, Irf1*, and *Isg15,* which have been implicated in β cell death in T1D models (38, 65, 66) (**Supplemental Figure S3C and S3D**). The β-cell population was comprised of 5 clusters (**Figure 6C**). Notably, the β cell cluster exhibiting the greatest reduction in relative abundance following E2 treatment was characterized by elevated expression of genes involved in antigen presentation (e.g., MHC I genes *H2-K1*, *H2-D1*, *B2m*) and interferon signaling (*Irf1, Stat1, Stat2*) (**Figure 6F**), suggesting a highly inflamed and immunogenic phenotype. Furthermore, E2 treatment in vivo reduced expression of several β-cell maturity markers, including *Slc2a2*, *Pdx1*, and *Nkx6-1* (**Figure 6G**), suggesting that E2 reduces the expression of both β-cell-specific antigens and antigen presentation machinery. This observation aligns with prior reports in the NOD model, wherein β cells with diminished expression of canonical identity genes exhibit increased survival during autoimmune attack (48). These findings suggest that E2 may protect β cells, at least in part, by downregulating features that increase their immune visibility. Collectively, these findings indicate that 4 weeks of E2 treatment attenuates β-cell immunogenicity by suppressing interferon signaling, chemokine production, antigen presentation, and markers of β-cell maturity—each of which may contribute to the reduced diabetes incidence observed in E2-treated mice in this and other studies.

### Estradiol treatment promotes tolerogenic phenotypes in islet-infiltrating immune cells

β cells play an active role in shaping the local immune environment. Prior studies demonstrated that dampening β-cell-intrinsic inflammation can drive less inflammatory phenotypes in islet-infiltrating immune cells (20, 21), suggesting a reciprocal interaction that may be influenced by E2 treatment. Pursuant to these observations, we next examined transcriptional immune phenotypes in the scRNAseq data. After 4 weeks of E2 treatment, the relative proportions of macrophages, T cells, and B cells were similar between treatment groups; although of marginal statistical significance (p=0.07), dendritic cells (DCs) were reduced in the E2 group (**Figure 7A**). Notably, the DC population expressed markers of plasmacytoid DCs (pDCs), including *Siglech*, *Bst2*, *Tcf4*, and *Tlr7* (**Figure 7B**). pDCs are antigen-presenting cells that produce IFN-α, and this population is overrepresented in the peripheral blood of individuals with recent-onset T1D (15, 33, 67).

**Figure 7:**
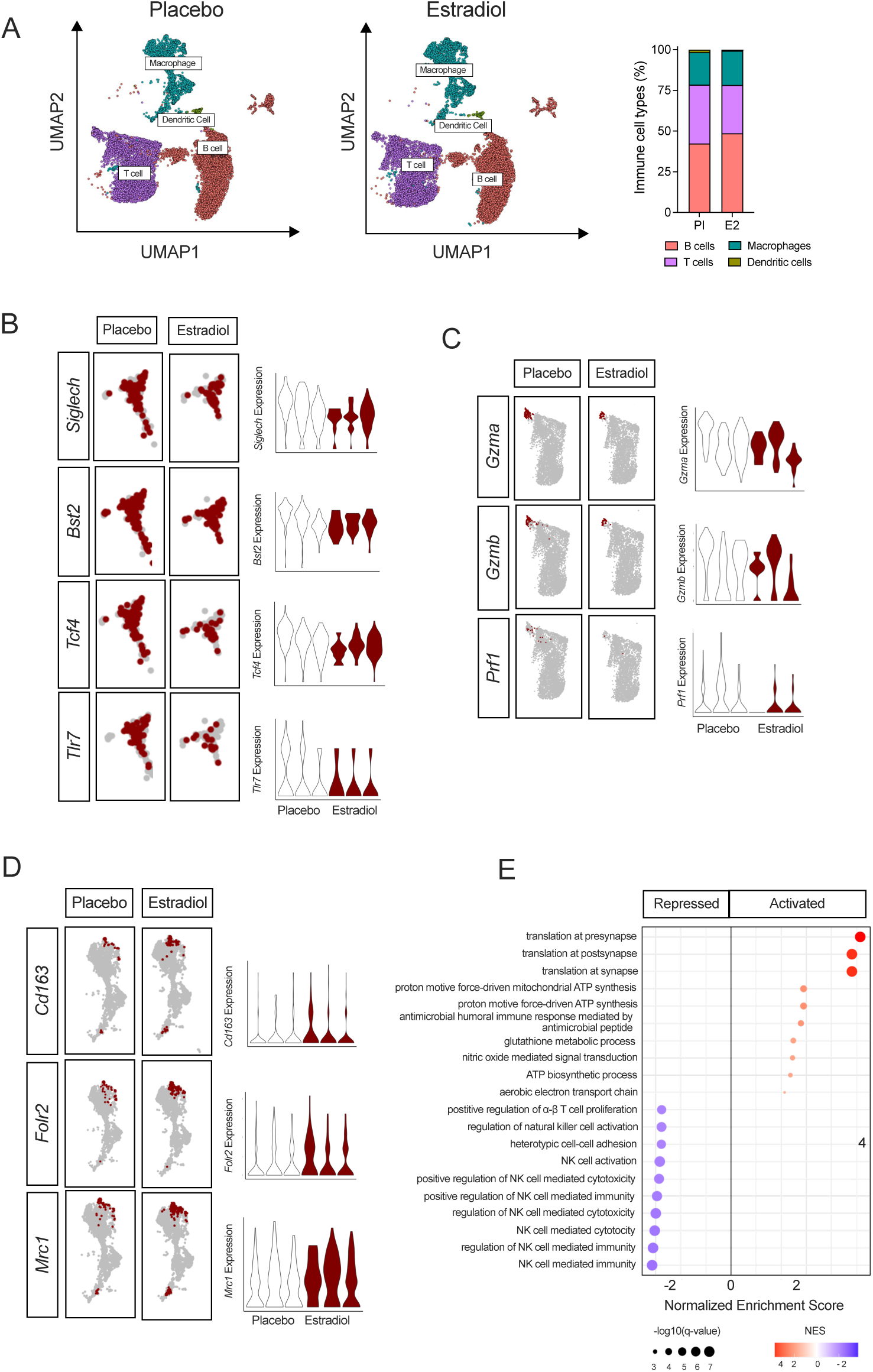
Single-cell RNA-seq revealed immune cell differences after estradiol treatment of female NOD mice. Prediabetic female NOD mice were treated with placebo or E2 pellet implants for 4 weeks and isolated islets were subjected to scRNA-seq. N=3 biological replicates per group. (**A**) UMAP embeddings of the immune cell clusters of islets isolated from mice treated with placebo (*left panel*) or E2 (*middle panel*). Percent of immune cells identified within the islet immune cell clusters (*right panel*). (**B**) Feature plots of *Siglech*, *Bst2*, *Tcf4,* and *Tlr7* gene expression in the dendritic cell clusters (*left panels*). Quantification of *Siglech*, *Bst2*, *Tcf4,* and *Tlr7* gene expression (*right panels*). (**C**) Feature plots of *Gzma*, *Gzmb1*, and *Prf1* gene expression in the T cell cluster 5 (*left panels*). Quantification of *Gzma*, *Gzmb1*, and *Prf1* gene expression (*right panels*). (**D**) Feature plots of *Cd163*, *Folr2*, and *Mrc1* gene expression in the macrophage cell clusters (*left panels*). Quantification of *Cd163*, *Folr2*, and *Mrc1* gene expression (*right panels*). (**E**) GSEA of the immune cell clusters.

Among the genes most differentially expressed in immune cells between E2 and placebo were markers of cytotoxic T effector cells, including cytolytic granzymes (*Gzma, Gzmb*) and perforin (*Prf1*); these were primarily expressed in a small cluster of *Cd8*-expressing T cells across treatment groups (**Figure 7C**). On the other hand, among the few genes more highly expressed in E2 immune cells than in placebo were *Cd163, Fol2r,* and *Mrc1,* each primarily expressed in macrophages and encoding markers of the anti-inflammatory M2-like phenotype (**Figure 7D**). GSEA of the immune cell compartment showed that pathways related to lymphocyte cytotoxicity, activation, and proliferation among many others, were suppressed with E2 treatment, while pathways enriched by E2 treatment related to mitochondrial ATP production and protein translation, as in the β cells (**Figure 7E**). Taken together, these findings are consistent with prior studies suggesting that β-cell-intrinsic suppression of proinflammatory, antigen presentation, and chemokine pathways can influence local immune phenotypes (20, 21), raising the possibility that the immune changes observed with E2 treatment may be secondary to upstream changes in β-cell signaling.

### Pretreatment of islets with E2 attenuates T-cell activation and proliferation in vitro and reduces hyperglycemia in vivo

Our findings in this study indicate that E2 suppresses IFN signaling in β cells and downregulates key markers of β-cell identity. To investigate whether these E2-driven changes at the islet alter islet immunogenicity, we utilized an islet cell/immune cell co-culture system. Islets isolated from NOD/SCID mice were dispersed to single cells then cultured for 48h in the presence of E2 or vehicle (**Figure 8A**). Islet cells were then co-cultured for 48 or 72 hours with splenocytes from BDC2.5 NOD mice, whose CD4+ T cells express a transgenic chromogranin A-reactive T-cell receptor (TCR) that targets islets (**Figure 8A**) (68). Positive controls for T cell activation included incubations with concanavalin A (ConA) and autoreactive BDC2.5 peptide. After 72 hours, all positive control incubations revealed activation of T cells (including T cell proliferation and IFN-γ secretion into media) by flow cytometry or ELISA (**Figure 8B-F**).

**Figure 8:**
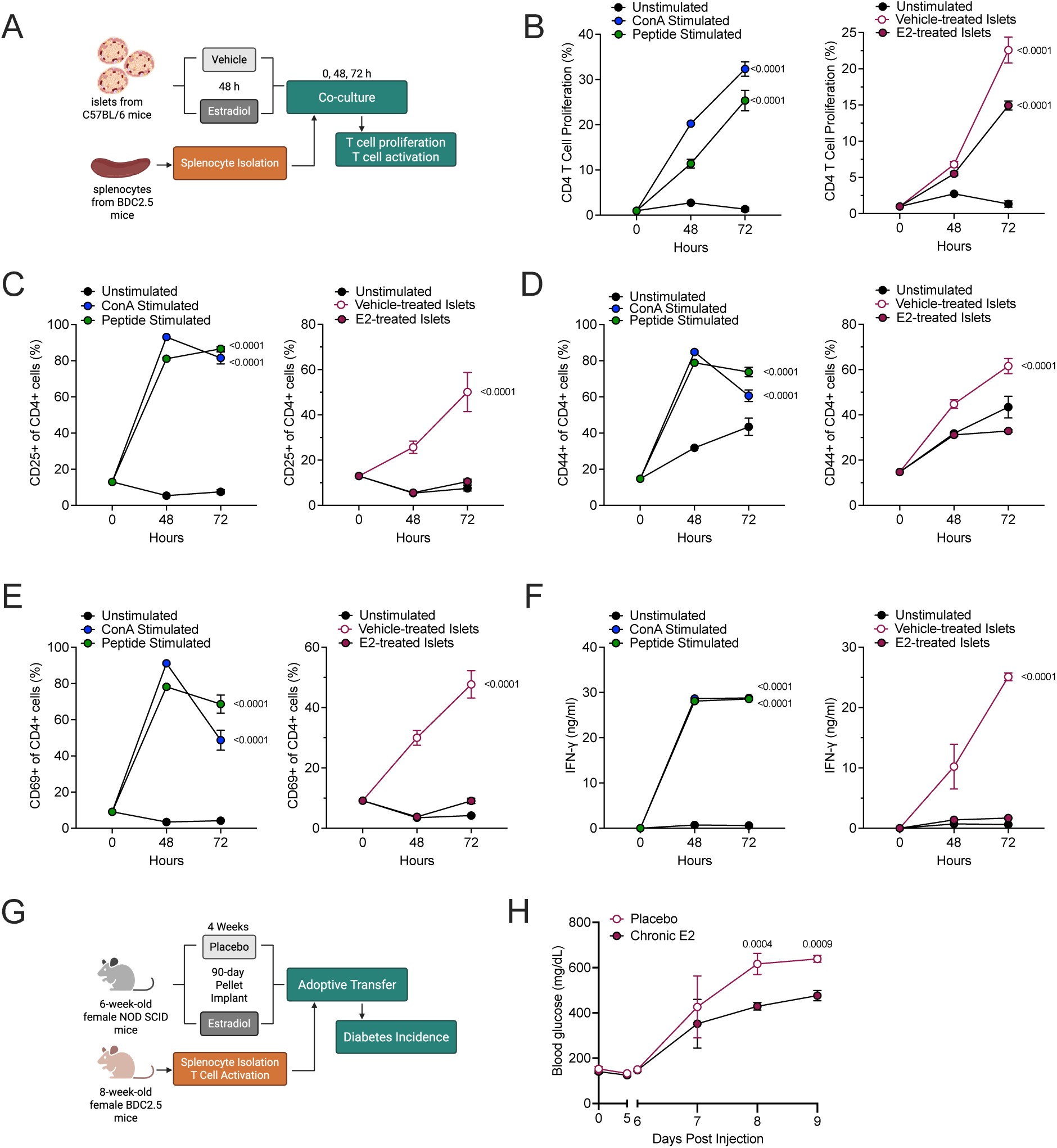
Estradiol treatment reduces islet immunogenicity and hyperglycemia. Splenocytes from BDC2.5 mice were co-cultured with C57BL/6 islet cells pre-treated with E2 or placebo. (**A**) Experimental design for co-culture studies. (**B**) CD4+ T Cell proliferation by CellTrace Violet, (**C**) quantification of CD25+ of CD4+ cells (%), (**D**) quantification of CD44+ of CD4+ cells (%), and (**E**) quantification of CD69+ of CD4+ (%) via flow cytometry, and (**F**) supernatant IFN-γ measured via ELISA, all performed on splenocytes stimulated with concanavalin A or BDC2.5 peptide (*left panels*) or stimulated by co-culture with islets pre-treated with E2 or placebo (*right panels*). N=3-6 biological replicates. (**G-H**) Female NOD SCID mice were treated with placebo or E2 90-day pellet implants prior to adoptive transfer of BDC2.5 splenocytes. (**G**) Experimental design. (**H**) Random-fed blood glucose values at day 8 post adoptive transfer. N=10 mice per group. Statistical tests: Two-way Anova with Dunnett’s post-hoc test: **B-F**; Unpaired t-test: **G**. Data are represented as mean ± SEM.

Similarly, co-cultures of vehicle-treated islets also elicited a robust T cell proliferative response and IFN-γ secretion (**Figure 8B-F**). By contrast, significantly reduced T cell proliferation (**Figure 8B**) and markers of T cell proliferation and activation (**Figure 8C-E**) with reduced IFN-γ secretion (**Figure 8F**) were observed in co-cultures with islets previously treated with E2 compared to co-cultures with vehicle-treated islets. These data suggest that E2 exerts direct effects on the islet that reduce their potential to drive immunogenicity, further underscoring an active role for the islet in shaping downstream immune responses.

To determine whether β-cell-intrinsic effects of E2 are sufficient to influence severity of disease outcomes, we employed an aggressive adoptive transfer model in which activated islet-reactive CD4⁺ T cells from BDC2.5 NOD mice were transferred into immunodeficient NOD/SCID mice—a setting of rapid, IFN-driven β-cell destruction (69, 70). Recipient mice were pretreated with placebo or E2 pellets from 6 to 10 weeks of age, followed by T cell transfer and monitoring of blood glucose levels (**Figure 8G**). As expected in this highly aggressive model, all mice developed overt diabetes (glucose >250 mg/dL) by 8 days post-transfer. However, E2-treated recipients exhibited significantly lower blood glucose levels compared to placebo-treated controls (**Figure 8H**). These findings suggest that E2 confers partial protection against β-cell dysfunction and loss, highlighting a likely direct effect on the β cell that limits the severity of hyperglycemia even in the face of robust autoimmune attack.

## DISCUSSION

T1D incidence has increased steadily over the decades, a trend largely attributed to the interplay between genetic predisposition and environmental triggers (71–74). A less widely recognized but consistent observation is the higher prevalence of T1D in males after puberty, with an approximate 1.5:1 male-to-female ratio (7, 8). Both of these phenomena may be rooted in the notion of β-cell “susceptibility”—the idea that intrinsic β-cell dysfunction, particularly under inflammatory stress, amplifies vulnerability to autoimmune attack and promotes disease progression (75–79). In this study, we leveraged the post-pubertal sex bias in T1D incidence to interrogate whether β-cell responses to inflammation differ by sex and contribute to disease risk. Our findings demonstrate that: (a) male human islets exhibit a more robust inflammatory response to cytokine exposure than female islets; (b) E2 suppresses islet inflammatory signaling and β-cell maturity markers in vitro and in vivo; (c) E2 treatment protects against autoimmune diabetes development in the NOD mouse model, due at least in part to islet-cell-intrinsic effects.

To model the early inflammatory milieu of T1D pathogenesis in vitro, we incubated human islets with a combination of IFN-γ and IL-1β, a cytokine cocktail (PIC) that has been widely used to mimic the proinflammatory microenvironment observed in preclinical diabetes (31, 32). This approach allowed us to both leverage publicly available transcriptomic and proteomic datasets using the same cytokine stimuli for discovery, and interrogate intrinsic islet responses to inflammation independent of infiltrating immune cells. Notably, our validation dataset—generated independently from different donors—recapitulated the sexually dimorphic features observed in the discovery cohort, underscoring the robustness of the findings. Male islets mounted a more extensive response to PIC, marked by a stronger induction of interferon-related pathways, particularly those associated with IFN-α signaling. Prior work has shown that IFN-α signaling promotes ER stress, neoantigens formation, HLA class I hyperexpression, and chemokine secretion (e.g., CXCL10), all of which enhance immunogenicity and attract immune cells (18, 19). Disruption of this pathway, as with deletion of HLA class I, allows human islets to evade immune surveillance when transplanted into immunocompetent mice (80). Proteomic analyses supported this observation: PIC-treated male islets showed a greater induction of interferon-stimulated proteins consistent with heightened β-cell immunogenicity. Accordingly, the enhanced IFN-α signaling in male islets may contribute to increased immune visibility and vulnerability.

In contrast, female islets showed enrichment in tissue development and remodeling pathways, including EMT and Wnt signaling. These processes have been associated with immune evasion in cancer and β-cell dedifferentiation, potentially rendering β cells less recognizable to autoreactive T cells (41, 42, 44, 50, 81–83). In this regard, β cells with reduced expression of identity markers and elevated PD-L1 are more likely to persist in autoimmune settings (47, 48, 84). These data raise the possibility that the female islet state confers a degree of immune privilege by limiting both recognition and destruction during inflammatory stress.

T1D incidence diverges between the sexes around puberty, with post-pubertal males showing increased susceptibility (7), implicating sex hormones as modulators of disease risk. While prior studies have established protective effects of 17-β estradiol (E2) in β cells exposed to oxidative, inflammatory, or metabolic stress (27–29, 85), the cellular and molecular basis for this protection has remained incompletely defined. Our study advances this understanding by demonstrating that female human islets exhibit a less proinflammatory response to cytokine stress and show enrichment for estrogen-responsive transcriptional programs. Furthermore, direct E2 treatment of human islets suppressed inflammatory signaling pathways—particularly those associated with interferon responses—that were more prominently activated in male islets. These observations not only align with prior murine studies showing protection of islets by estrogens in models of T1D (85, 86) but also extend those findings by bridging them to human β-cell biology and the sex disparity in human T1D incidence. Importantly, our transcriptomic and proteomic data together suggest that E2 reprograms β cells toward a state that is both less immunogenic and more resilient to inflammatory insult, and this notion is further bolstered by the diminished T cell activation and proliferation that was stimulated by exposure to islets pre-treated with E2 vs. control. These data could help to explain, at least in part, the female bias in T1D resistance.

Our findings present important opportunities for future study, particularly regarding the specific estrogen receptor pathways that mediate the restrained inflammatory response in female islets and the protective effects of E2 against diabetes in the NOD model. Earlier studies demonstrate that knockout of *Esr1* (encoding the receptor ER-α) or of *Gper1* (encoding a membrane estrogen receptor) each sensitizes female mice to streptozotocin-induced β-cell death and diabetes (27, 87). In our validation cohort, incubation of human islets with PIC induced a decrease in *Esr1* in male islets that was not observed in female islets, suggesting a possible role for ER-α in the sexually dimorphic PIC response. A potential role for ER-β cannot be ruled out, although studies suggest only a minor contribution of the receptor to β-cell cytoprotection (87). While the present study shows that E2 stimulates developmental pathway signaling in vitro and decreases markers of β-cell maturity in vivo, other studies suggest that signaling through GPER1 can maintain or restore β-cell maturity (88, 89), underlining the diverse roles of E2 signaling through different receptors or pathways. Future studies using receptor-selective agonists or conditional receptor deletions in the NOD model could clarify these mechanisms.

Some limitations of our study should be noted. While our findings support a protective role for E2 at the level of the islet, we cannot explicitly exclude a role for E2 in the immune compartment. It remains possible that direct actions of E2 on innate and adaptive immune cells may also contribute to disease protection, and future studies using cell-specific genetic perturbations will be necessary to disentangle these effects. Second, although we focused on estradiol as the primary mediator of female-biased protection, additional sex-specific factors— such as androgen signaling, sex chromosome gene dosage, and microRNA expression—may also influence islet inflammation and autoimmunity and warrant further investigation. Notably, androgens have also demonstrated protective anti-inflammatory effects both in the NOD mouse and in murine models of autoimmunity (90, 91). Finally, human islet studies are inherently subject to donor variability, which may confound the interpretation of the results. However, the reproducibility of our findings across independently derived transcriptomic datasets, with support from proteomics, from different donors, lends confidence to the observed sex differences and supports their relevance to human disease.

Taken together, our findings provide mechanistic insight into how sex differences, mediated in part through estradiol signaling, shape islet-intrinsic inflammatory responses that may influence the course of autoimmune diabetes. By integrating transcriptomic and proteomic data from human islets in vitro with single-cell analyses in NOD mice in vivo, we demonstrate that estradiol reprograms β cells away from immunogenic and mature phenotypes toward a state that may enhance resilience in the face of inflammatory stress. While the complexity of sex differences in T1D extends beyond estradiol signaling, our data highlight the therapeutic potential of modulating β-cell states to restore immune tolerance and preserve islet function.

Recognizing the molecular determinants of β-cell susceptibility offers a pathway toward precision prevention strategies that are responsive to sex and other individual variables.

## METHODS

### Sex as a biological variable

Both male and female human islets were used for the study. Both male and female NOD mice were used for diabetes incidence and pancreatic tissue analysis. Single cell RNA sequencing was performed on female NOD islets only. For adoptive transfer experiments, the recipient NOD/SCID mice were female.

### Human islet culture

De-identified human islets from non-diabetic cadaveric donors were provided by the Integrated Islet Distribution Program (IIDP) and the University of Alberta Diabetes Institute Islet Core (Edmonton, Alberta, Canada) (**Table 1** and **Supplemental Table S1)**. Culture medium used for human islet experiments was Prodo Islet Media (Standard) supplemented with human AB serum, glutamine and glutathione (Prodo), and ciprofloxacin (Fisher). Islets were cultured in the presence or absence of a proinflammatory cytokine (PIC) cocktail containing 1000 IU/mL human IFN-γ (R&D Systems; 285-IF-100) and 50 IU/mL human IL-1β (R&D Systems; 201-LB-005), or with 10 nM E2 (Millipore Sigma, E2758) for 18 hours.

### Bulk RNA sequencing

Human islet lysates were stored in RLT plus buffer containing 1% β-mercaptoethanol, and RNA was extracted using the RNeasy Plus Micro kit (Qiagen) according to the manufacturer’s instructions. Samples were submitted to the University of Chicago Genomics Core for sequencing using the NovaSeq-X system (Illumina). Reads were aligned to the *Homo sapiens* genome build hg38 and quantified using Salmon. Transcripts were summarized to the gene level using tximport (v1.30.0). Owing to high variability between human samples, raw counts were input into ComBat-Seq (92) for batch correction using default parameters. The batch corrected matrix, DESeq2 was then analyzed using DESeq2 (v1.42.1) in R. Counts normalization was conducted via variance stabilization transformation (vst), and the normalized counts were used for principal component analysis and heatmap visualizations. Genes with raw counts greater than ten in at least ten samples were kept for differential expression testing, and the Wald statistic was applied to rank genes for gene set enrichment analysis. The Benjamini-Hochberg method was used for p-value adjustment, and genes with adjusted p-values < 0.05 and an absolute fold change >1.5 were considered differentially expressed. Pathway analysis was performed using Gene Ontology (GO).

### Reanalysis of RNA-seq datasets

Re-analyses of a previously published dataset available via the NCBI’s Gene Expression Omnibus (<u>GSE169221</u>) was performed using ROSALIND®, with a HyperScale architecture developed by ROSALIND, Inc. (San Diego, CA). Reads were trimmed using cutadapt (93). Quality scores were assessed using FastQC. Reads were aligned to the Homo sapiens genome build hg38 using STAR (94). DEseq2 was used to calculate fold changes and p-values (95). Overrepresentation analyses were performed using Enrichr (96–98). Gene set enrichment analysis was performed using the GSEA 4.3.2 software (99).

FASTQ files of 10x Genomics scRNAseq data for human islets from nondiabetic, AAB+, and T1D donors were downloaded from the data portal of the HPAP (https://hpap.pmacs.upenn.edu) (52). Cell Ranger version 6.1.2 was used to perform quality control and align reads to human genome build GRCh38 (100). SoupX version 1.6.1 (101) was used to remove ambient mRNA, and doublets were removed with scDblFinder version 3.16 (102). Cells retained for further analysis were those with 200-9,000 detected features, <25% mitochondrial reads, and fewer than 10,000 counts. Seurat function SCTransform (103, 104) was used to normalize counts, removing effects of library depth and regressing out the variation from the mitochondrial reads’ ratio. Principal component analysis was performed using the top 3,000 variable genes, and Harmony 0.1.1 (105) was used to integrate all samples based on the top 50 principal components, setting donor identity and reagent kits as primary confounders.

Human islet cell types were annotated using scScorter version 0.0.2 (106). A total of 10,167 β cells remained after all above steps, including 4,730 cells from nondiabetic donors, 2,372 cells from single-autoantibody–positive donors, 2,480 cells from double-autoantibody–positive donors, and 585 cells from donors with T1D. GSEA was performed in R using the msigbdr package (https://igordot.github.io/msigdbr/) using the Hallmarks Gene Sets.

### Mass spectrometry-based proteomics

Human islet lysates were stored in RLT plus buffer containing 1% β-mercaptoethanol, and acetone-based protein elution was performed on the flow-through from the RNeasy MinElute spin column from the RNeasy Plus Micro kit (Qiagen) according to manufacturer instructions. A data-independent acquisition approach was used for the quantitative proteomics analysis using liquid chromatography with tandem mass spectrometry (LC-MS/MS) on QExactive HF mass spectrometer (Thermo Fisher Scientific). (107). Acetone precipitated proteinwas suspended in 8 M urea, 50 mM Tris-HCl (pH 8.0), 75 mM NaCl, and complete protease inhibitor cocktail (Sigma 5892791001) buffer. Disulfide bonds were reduced by adding dithiothreitol (5 mM, final concentration), followed by an alkylation step using iodoacetamide (40 mM, final concentration). Samples were diluted 4-fold with 50 mM tris HCl (pH 8.0) and digested with an 1/50 (enzyme/protein) ratio of sequencing grade endoproteinase LysC (VWR) for 2 h at 25°C, followed by sequencing grade trypsin (Promega) overnight at 25°C. The reaction was quenched using trifluoroacetic acid (1%, final concentration). Samples were desalted using 15-mg C18-E Strata 96-well plate, and peptide concentrations were estimated with a BCA assay.

Five hundred nanograms of peptides were separated in a C18 column (70 cm 3 75 mm i.d., Phenomenex Jupiter, 3 mm particle size, 300 Å pore size) connected to an Acquity M-Class Nano UHPLC system (Waters). Data were collected in data-independent acquisition with tandem mass spectra from the range of 400–900 m/z with 10 m/z increment windows (10.0 m/z isolation width; 30% normalized collision energy; 70,000 resolution at 400 m/z). Post-LC-MS/ MS, an *in-silico* library was created using the human Swissprot database (September 12, 2024, 20,656 sequences) on our inhouse DMS data analysis interface using DiaNN. Only fully tryptic peptides were considered with up to two missed cleavages. Carbamidomethylation of cysteine was set as a fixed modification; acetylation of protein N-terminus and oxidation of methionine residues were set as variable modifications. Precursor peptide length was set to a range of 7-30, m/z 400-900 and charge 2-4. The final library contained peptides filtered with 1% false discovery rate (FDR) at protein, peptide, and peptide/spectrum match levels. The DIA sample runs were processed in the same workflow matching against the generated Insilco library, using an FDR of 1%.

Data was evaluated for potential sample outliers using a robust Mahalanobis distance (RMD) measure based on correlation with samples from the same treatment group, median absolute deviation (MAD), proportion of observations missing, and skewness of sample profiles (108). Sparse proteins without enough data to conduct a statistical test were filtered from the dataset. A protein was required to have a non-missing value for the control treatment and at least one other treatment, within the same donor, for at least two donors. Data was normalized via median centering. Comparisons of mean protein and gene expression of each group back to the control group were performed. Datasets were analyzed with all samples and also broken into separate datasets for males and females only and analyzed for these subsets. A generalized linear model with treatment group as a fixed effect and donor as a random effect was fit to each protein using the *lme4* package in R (109). The mean for each treatment group was compared to the control group for differences in mean abundance, and a Holm multiple comparison adjustment was conducted (110). Enrichr was used to perform overrepresentation analyses (96–98).

### Animals and procedures

All mice were housed in a specific-pathogen-free facility at the University of Chicago with ad libitum access to food and water. NOD/ShiLtJ (NOD; JAX #001976), NOD.Cg-Tg(TcraBDC2.5,TcrbBDC2.5)1Doi/DoiJ (NOD.BDC2.5; JAX #04460), NOD.Cg-*Prkdc^scid^*/J (NOD/SCID; JAX #001303), and C57BL/6J (JAX #000664) mice were purchased (Jackson Laboratories).

Six-week-old male and female NOD mice were implanted subcutaneously with a 90-day, 0.5 mg pellet containing E2 or matched placebo (Innovative Research of America, NE-121, NC-111). For diabetes incidence experiments, blood glucose was measured up to twice weekly via tail vein using an AlphaTrak glucometer beginning at 10 weeks of age. Mice were euthanized after diabetes incidence (two consecutive readings > 250 mg/dL) or at 25 weeks of age, whichever came first. For insulitis, β cell mass, and scRNAseq studies, mice were euthanized at 10 weeks of age. Serum and tissues were collected upon euthanasia at the conclusion of each study.

For adoptive transfer experiments, recipient 6-week-old NOD/SCID females were implanted with a 21-day, 0.1 mg pellet containing E2 or matched placebo (Innovative Research of America). A single-cell suspension of splenocytes was prepared from 9-week-old female NOD.BDC2.5 mice. From splenocytes, CD4+ T cells were purified and activated in 6-well plates with αCD3/αCD28 beads (Gibco) and 100U/mL recombinant human IL-2 (Peprotech) for 72h.

CD4+ T cells were then expanded into T-75 flasks with 100U/mL IL-2 for 72h, washed with PBS, and diluted to 1 x 10^7^ cells/mL in PBS. Recipient NOD SCID females were injected intraperitoneally with 1 x 10^6^ CD4+ T cells each at 10 weeks of age, 1 week after pellet washout, and blood glucose was monitored via tail vein.

### Islet isolation and mouse cell culture

Mouse islets were isolated as described (111). Briefly, collagenase was injected into the pancreatic bile duct to inflate the pancreas, which was then removed and dissociated via rocking and syringe mixing in a 50 mL tube. Dissociated tissues were then placed within a gradient of Histopaque and HBSS and centrifuged at 2080 rpm for 18 minutes. After passing the supernatant through a 70-uμmfilter, the islets were rinsed into a culture plate in RPMI medium containing 10% fetal bovine serum and 1% penicillin/streptomycin. Healthy islets were handpicked under microscopy and transferred to fresh media, after which they were allowed to recover at 37°C and 5% CO_2_ overnight before experimentation.

Murine β cell line MIN6 (112) was cultured in high glucose Dulbecco’s Modified Eagle’s Medium (DMEM) containing 15% fetal bovine serum, 1% penicillin/streptomycin, and 1% L-glutamine. MIN6 cells and mouse islets were treated with a proinflammatory cytokine cocktail containing 25 ng/mL mouse IL-1β (R&D Systems; 401-ML-010), 50 ng/mL mouse TNF-α (R&D Systems; 410-MT-010), and 100 ng/mL mouse IFN-γ (R&D Systems; 485-MI-100) for 18 hours, or with 17-β estradiol (E2) (Millipore Sigma, E2758) (concentration and duration detailed in each experiment in Results). E2 was dissolved in EtOH, and the final EtOH concentration in all E2 cultures and corresponding controls was 0.1%.

### Islets and BDC2.5 T cells co-culture assay

Islets were isolated from (N=6) NOD-SCID mice (NOD.Cg-Prkdcscid/J; Strain 001303) following standard protocols (111) and kept overnight in islet media for stress relief. Approximately 80 intact islets/well were cultured in the presence of E2 (10 μM/ml) or EtOH as vehicle for 48 h. After incubation, islets were washed with PBS and dissociated using 500 μL Accutase (Fisher Scientific, NC9839010) in a 37 °C water bath for 30 min with intermittent pipetting. Dissociated single islets cells were counted and used for coculture. Splenocytes were harvested from

NOD.Cg-Tg(TcraBDC2.5) mice (Strain 004460). Briefly, spleens were processed through a 70 μm strainer (Corning, CLS352350), subjected to RBC lysis (Sigma Aldrich, R7757), washed twice by PBS, and finally resuspended into complete DMEM for counting. Cells were stained with Cell Trace Violet (Thermo Fisher, Invitrogen) per manufacturer’s protocol. For co-culture, 5×10⁵ splenocytes were incubated with 8×10⁴ single islet cells in complete DMEM at 37 °C, 5% CO₂ for 48 or 72 h. Positive controls included stimulation with BDC2.5 peptide (0.05 μM; 2.5HIP: RGG-LQTLALWSRMD-GGR) or Concanavalin-A (2.5 μg/mL; Sigma, C2272). Each condition was performed in triplicate in two independent experiments. Supernatants were collected for measuring mouse-IFNγ by ELISA, and splenocytes were analyzed by flow cytometry for immune profiling.

### Flow cytometry

Flow cytometry was performed to evaluate T cell proliferation and activation marker expression. CTV-stained splenocytes from co-cultures were washed in FACS buffer (1% BSA in PBS) and stained with anti-mouse FITC-CD45, PerCP-Cy5.5-CD4, BV605-CD44, PerCP-Cy7-CD69, and PE-CD25 (BioLegend), followed by eFluor780™ fixable viability dye (65086514, Thermo Fisher Scientific). Single-color compensations were prepared with UltraComp eBeads Plus (Invitrogen, 01-3333-42). Cells were sequentially gated on FSC/SSC, live cells, singlets, CD45, and CD4. CD4⁺ T cells were analyzed for CTV dilution for proliferation and CD69, CD44, CD25 activation marker expressions. FMO controls were included for each fluorochrome for accurate gating. Sample were acquire using Attune NxT Flow Cytometer, and data was analyzed by using FlowJo™ v10 (BD Biosciences).

### Enzyme-linked immunosorbent assay (ELISA)

Mouse IFN-γ ELISA was performed on co-culture supernatant. 96-well plates (Corning, 3361) were coated overnight at 4 °C with 25 ng/ml monoclonal rat anti-murine IFN-γ (BD Bioscience, #551216) diluted in carbonate coating buffer pH 9.6. Mouse IFN-γ ELISA antibody pairs were purchased from BD Biosciences, and assay was performed according to manufacturer’s protocol. ELISAs were read on SpectraMax-iD5 and analyzed using SoftMax-Pro. 17-β estradiol (E2) was measured in mouse serum using an estradiol ELISA kit (Cayman, #501890) according to the manufacturer’s instructions. Before measurement, samples were subjected to methanol extraction and dried under inert gas according to the manufacturer’s instructions.

### Quantitative RT-PCR

RNA was extracted from samples stored in Qiagen RLT plus buffer containing 1% β-mercaptoethanol using the RNeasy Mini Kit. For cDNA synthesis, High-Capacity cDNA Reverse Transcription Kit (Applied Biosystems) was used according to manufacturer’s instructions.

Taqman primer-probes (Applied Biosystems) were used to detect desired mRNA targets via quantitative PCR using Bio-Rad CFX Opus. Relative gene expression was calculated using the comparative threshold cycle value (CT). Data are displayed normalized to the expression of *Actb* and then to the average of the control group expression (ΔΔCT).

### β-cell mass and insulitis quantification

Pancreata were fixed in 4% paraformaldehyde and embedded in paraffin, then serial 5 μm sections were cut from 3 levels (100 μm apart). 3 sections per mouse (1 from each level) were stained with anti-insulin primary antibody (ProteinTech, 15848-1-AP at 1:200 dilution). Label was developed using ImmPRESS peroxidase-conjugated secondary antibody and DAB peroxidase substrate (Vector Laboratories), followed by hematoxylin counterstaining. Slides were imaged using the Keyence BZ-X810 fluorescence microscope system. BZ-X800 Analyzer was used to quantify insulin positive area and total pancreas area. β cell mass was calculated by multiplying (insulin positive area/total pancreas area) by pancreas weight measured at the time of euthanasia. Insulitis scoring was performed by assessing the extent of immune cell infiltration at each islet: 1 = no immune infiltrates, 2 = infiltrates cover <50% of area, 3= >50% area covered by infiltrates, 4= infiltration over 100% of islet.

### Single-cell RNA sequencing

Isolated NOD mouse islets were dissociated into single cells by incubating in Accutase containing DNase and RNasin (Promega) for 8 minutes at 37°C and washed twice with 1% bovine serum albumin HBSS. Single cell suspension was submitted to the University of Chicago Genomics Facility for 10x library generation using Chromium Single Cell 3’ v 3.1, followed by sequencing on Illumina NovaSeq-X. Cell Ranger was used to align reads to the *Mus musculus* genome build GRCm39, count unique molecular identifiers (UMIs), call cell barcodes, and perform default clustering. Ambient mRNA was removed using SoupX, version 1.6.1, and clustering was performed utilizing Seurat version 5.1.0 in R version 4.3.1. Cells were filtered to those with at least 500 unique features and <5% mitochondrial genes. Replicates from each treatment group (estrogen vs. placebo, n=3 each) were merged using SCTransform, and integration was performed using Harmony version 1.2.3. Clustering was performed using Seurat and visualized by UMAP, and cluster markers were identified using FindAllMarkers (Wilcoxon’s rank-sum test) in Seurat. Differential gene expression was analyzed using FindMarkers. GSEA was performed on the endocrine and immune cell subsets using the ClusterProfiler package, version 4.10.1. Data visualizations were created using Seurat and ggplot2 version 3.5.1.

### Statistical Analyses

For comparisons involving 2 conditions, a 2-tailed student’s unpaired t-test was performed. A Mantel-Cox log-rank test was performed to compare diabetes incidence between treatment groups. GraphPad Prism, version 10, was used for these statistical analyses and corresponding visualizations. Statistical significance was defined as *P* < 0.05. Statistics used for analyses of RNAseq and proteomic analyses are described in more detail above.

### Study approval

The use of de-identified human samples was approved by the Institutional Review Board at the University of Chicago and considered exempt from human subjects research. Studies involving mice were performed under a protocol approved by the University of Chicago Institutional Animal Care and Use Committee.

### Data availability

RNA sequencing data will be uploaded to the public repository GEO and proteomics data will be uploaded to MassIVE upon acceptance. Any additional information required to reanalyze the data reported in this paper is available from the corresponding author upon request.

## ACKNOWLEDGEMENTS

This work was supported in part by National Institutes of Health grants R01 DK060581 and R01 DK124906 (both to RGM), U01 DK127786 (to RGM, CEM, ESN, and BJMWR), R01 DK135832 (to SAT), T32 DK131958 and T32 GM150375 (both to KLW), and T32 AI153020 (to JRE); a Breakthrough T1D postdoctoral fellowship (to JRE). This study utilized Diabetes Center core resources supported by National Institutes of Health grant P30 DK020595 (to the University of Chicago) and P30 DK097512 (to Indiana University) and utilized services of the University of Chicago Histology and Genomics Cores. We thank Chelsea M. Hutchinson Bunch at Pacific Northwest National Laboratory for assistance in the preparation of samples for proteomics, and Trinity-Paris Foster, Destini Wiseman, and Xiaxia Saavedra at the University of Chicago for assistance in adoptive transfer studies.

## AUTHOR CONTRIBUTIONS

KLW, ESN, WW, CEM, SRH, SAT, and RGM conceptualized the research; KLW, AAP, JRE, SS, BJMWR, LMB, SR, KF, and SP performed the investigations; SRH, HMT, JDP, RGM, and SAT provided project supervision; KLW, SAT, and RGM wrote the original draft; all authors contributed to discussions, edited the manuscript, and approved the final version of the manuscript.

---

**Supplemental Table S1:** Individual islet donor characteristics for the discovery and validation datasets.

| Donor ID | Source | Set | Omics | Age | Sex | BMI | HbA1c (%) | Islet Purity (%) |
| --- | --- | --- | --- | --- | --- | --- | --- | --- |
| SAMN08769390 | IIDP | Discovery | Transcript | 33 | Female | 34.2 | 5.5 | 95 |
| SAMN08769199 | IIDP | Discovery | Transcript | 43 | Female | 29.5 | 5.7 | 80 |
| SAMN08769206 | IIDP | Discovery | Transcript | 52 | Female | 25.9 | 5.6 | 90 |
| SAMN08769129 | IIDP | Discovery | Transcript | 44 | Female | 34.5 | N/A | 90 |
| SAMN08769191 | IIDP | Discovery | Transcript | 35 | Male | 24.2 | 5.4 | 95 |
| SAMN08769188 | IIDP | Discovery | Transcript | 30 | Male | 35.9 | 5.4 | 90 |
| SAMN08769132 | IIDP | Discovery | Transcript | 23 | Male | 24.8 | 5.3 | 95 |
| SAMN08769121 | IIDP | Discovery | Transcript | 28 | Male | 24.6 | 5.1 | 95 |
| SAMN08769124 | IIDP | Discovery | Transcript | 15 | Male | 24.6 | 5.1 | 80 |
| SAMN08769073 | IIDP | Discovery | Transcript | 28 | Male | 30.7 | 4.9 | 95 |
| SAMN34356133 | IIDP | Validation | Transcript/Protein | 20 | Male | 38.5 | 5.5 | 85 |
| SAMN34997905 | ADI | Validation | Transcript/Protein | 40 | Male | 30.4 | 5.4 | 95 |
| SAMN35848421 | IIDP | Validation | Transcript/Protein | 22 | Male | 31.79 | 5.1 | 90 |
| SAMN36704819 | IIDP | Validation | Transcript/Protein | 52 | Male | 25.9 | 4.9 | 85 |
| SAMN37529025 | IIDP | Validation | Transcript/Protein | 64 | Male | 23 | 4.8 | 90 |
| SAMN37638596 | IIDP | Validation | Transcript/Protein | 34 | Male | 27.2 | 5.5 | 85 |
| SAMN37844810 | ADI | Validation | Transcript/Protein | 63 | Male | 34.7 | 5.9 | 90 |
| SAMN37927922 | ADI | Validation | Transcript/Protein | 57 | Male | 26.7 | N/A | 95 |
| SAMN38117428 | IIDP | Validation | Transcript/Protein | 47 | Male | 28.1 | 5.3 | 95 |
| SAMN38750927 | IIDP | Validation | Transcript/Protein | 34 | Male | 24.4 | 5.4 | 95 |
| SAMN35985448 | ADI | Validation | Transcript/Protein | 68 | Female | 22.3 | 5.6 | 95 |
| SAMN36458803 | ADI | Validation | Transcript/Protein | 49 | Female | 28.5 | 5.2 | 80 |
| SAMN36823227 | IIDP | Validation | Transcript/Protein | 41 | Female | 38.4 | 5.2 | 75 |
| SAMN37217050 | ADI | Validation | Transcript/Protein | 53 | Female | 33.1 | 5.4 | 80 |
| SAMN37283209 | ADI | Validation | Transcript/Protein | 59 | Female | 19.9 | 5.3 | 85 |
| SAMN38143287 | ADI | Validation | Transcript/Protein | 41 | Female | 27.3 | 5.7 | 90 |
| SAMN38226661 | IIDP | Validation | Transcript/Protein | 37 | Female | 30 | 5.1 | 85 |
| SAMN38637296 | ADI | Validation | Transcript/Protein | 29 | Female | 31 | 5.1 | 70 |
| SAMN39461369 | IIDP | Validation | Transcript/Protein | 53 | Female | 24.7 | 5.6 | 90 |
| SAMN40160994 | IIDP | Validation | Transcript/Protein | 34 | Female | 22.5 | 5.1 | 80 |

**Supplemental Figure 1 (related to Figure 1):**
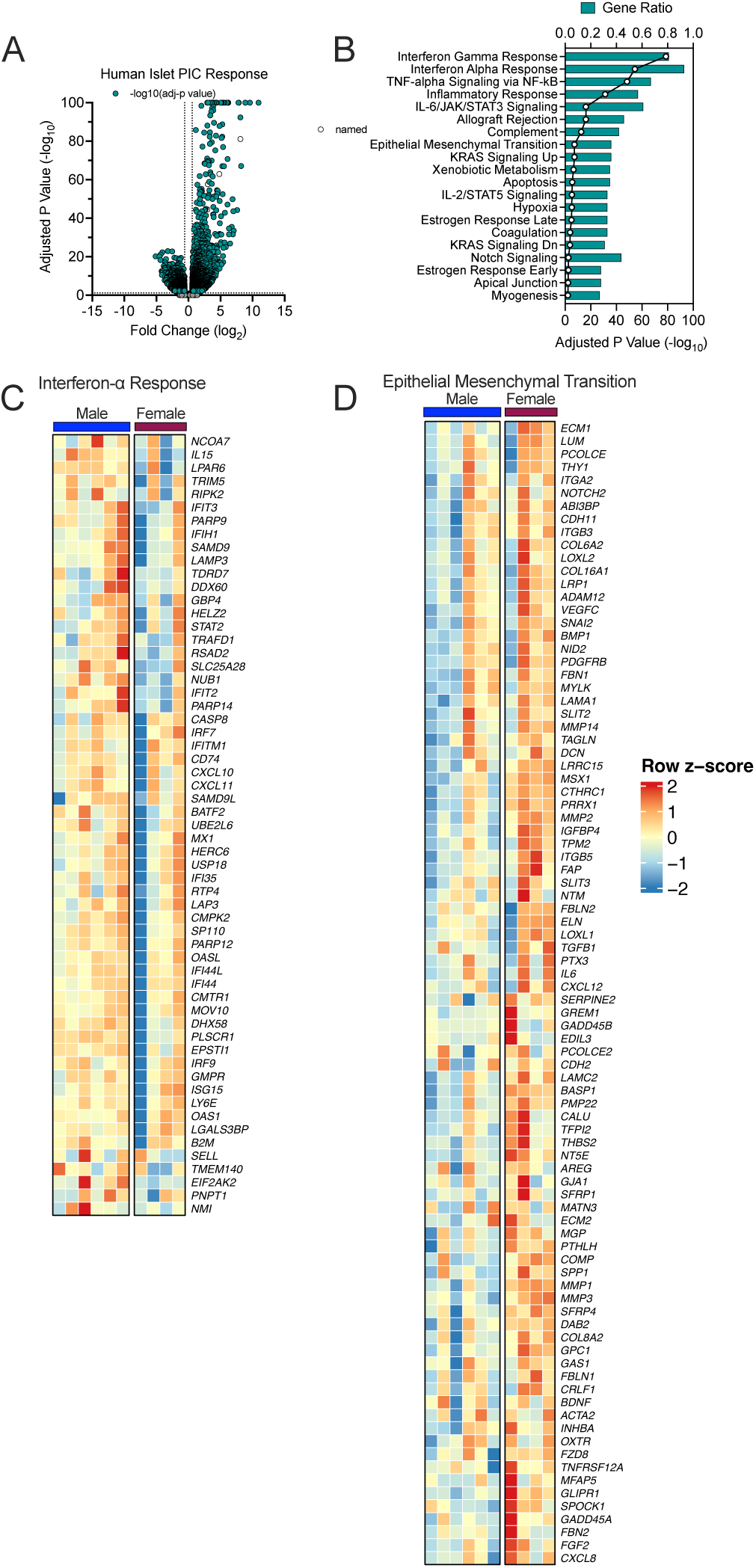
(***A***) Volcano plot of differentially expressed genes in male and female islets combined treated with PIC compared to vehicle. (***B***) Hallmark pathway analysis of differentially expressed genes in male and female islets combined treated with PIC compared to vehicle. (***C***) Heatmap of the leading edge genes from GSEA of Hallmark: Interferon Alpha Signaling in islets from male donors treated with PIC compared to female islets from female donors treated with PIC. (***D***) Heatmap of the leading edge genes from GSEA of Hallmark: Epithelial Mesenchymal Transition in islets from male donors treated with PIC compared to female islets from female donors treated with PIC. Islets from human donors: N=6 males and N=4 females.

**Supplemental Figure 2 (related to Figure 2):**
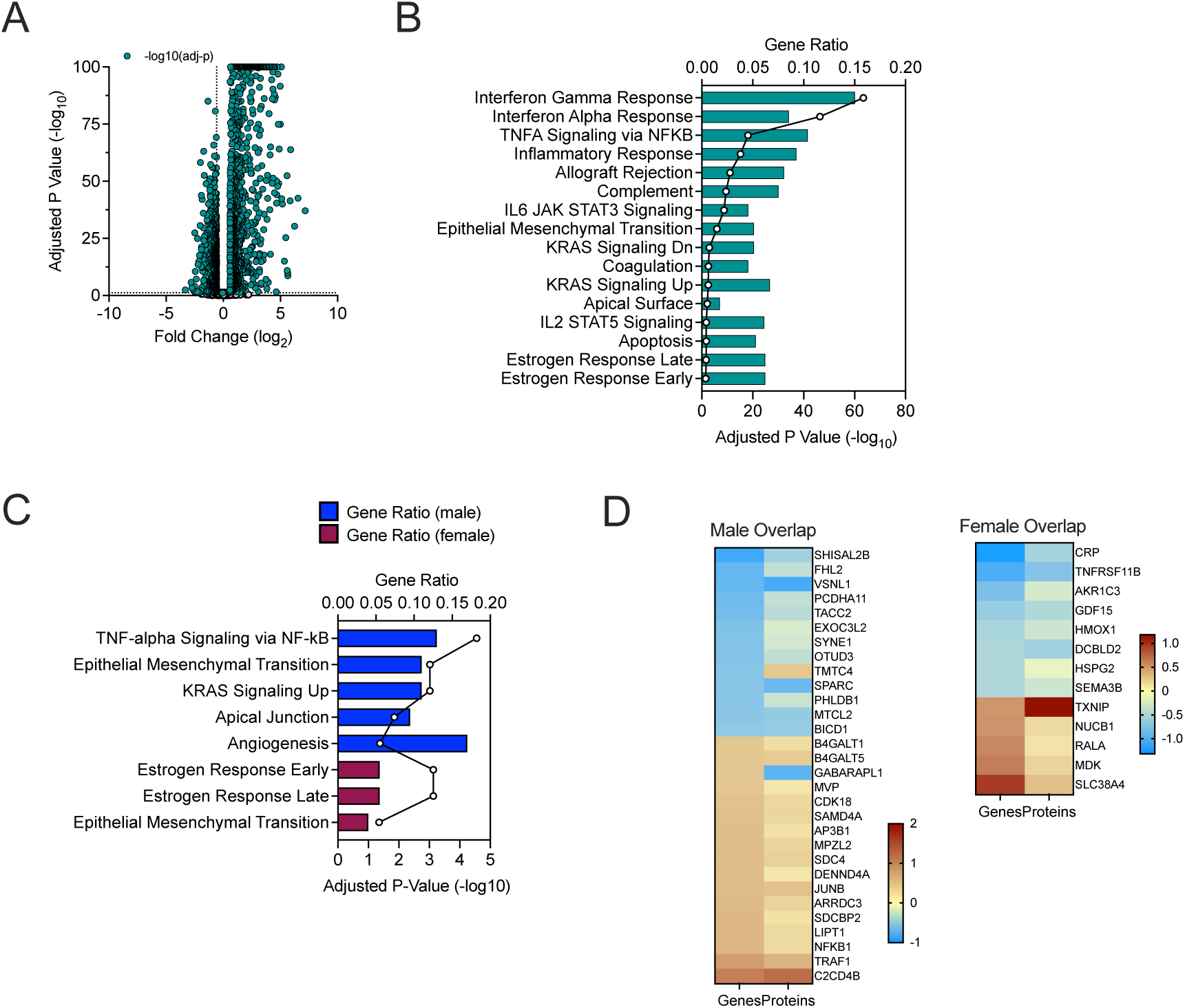
(***A***) Volcano plot of differentially expressed genes in male and female islets combined treated with PIC compared to vehicle. (***B***) Hallmark pathway analysis of differentially expressed genes in male and female islets combined treated with PIC compared to vehicle. (***C***) Hallmark pathway analysis of sex-specific differentially expressed genes. (***D***) Heatmap depicting overlap of male-specific (***left panel***) and female-specific (***right panel***) differentially expressed genes (DEG) and differentially abundant proteins (DAP) in response to PIC. Colors scaled by fold-change (log2) with PIC. Islets from human donors: N=10 males and N=10 females.

**Supplemental Figure 3 (related to Figure 6 and 7):**
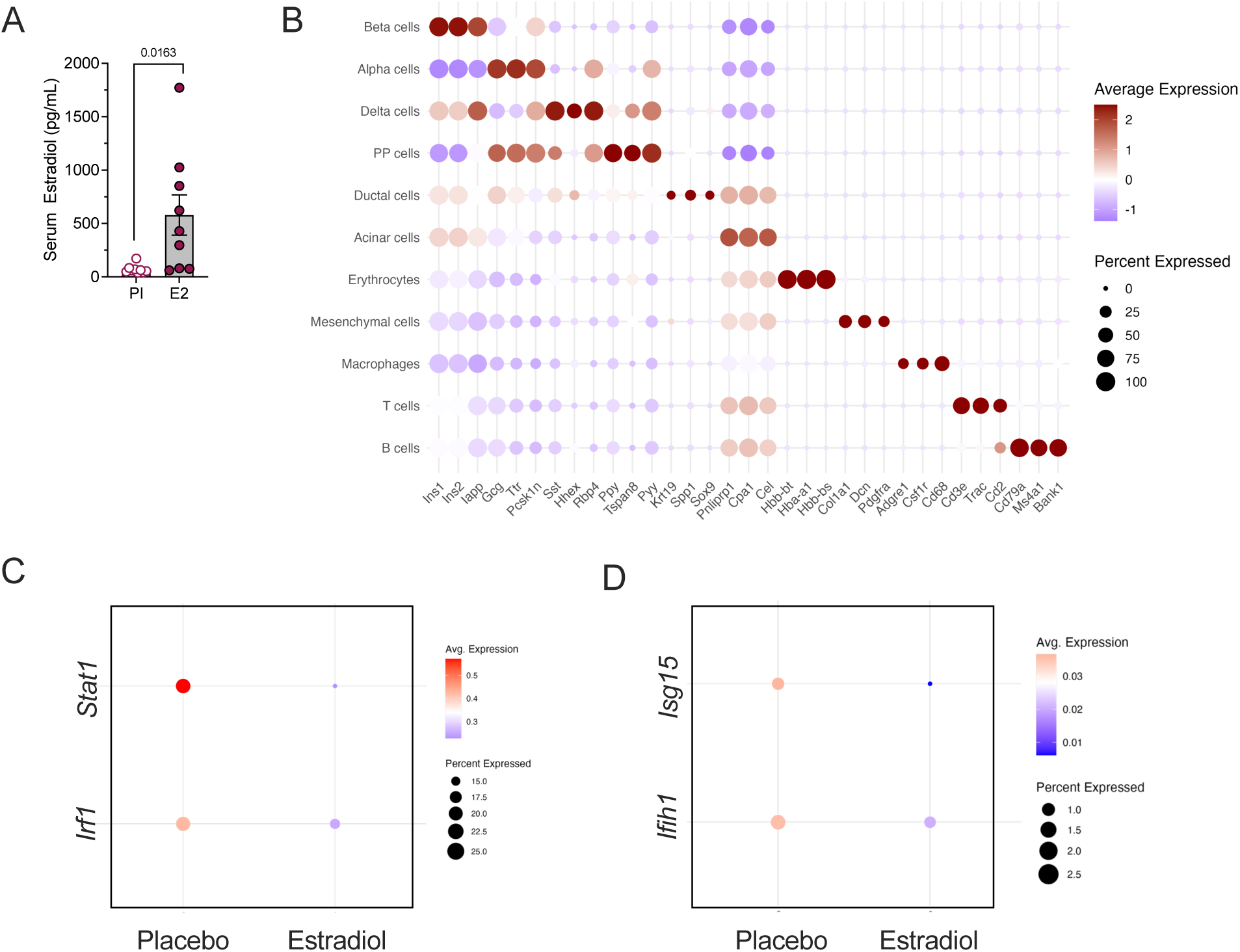
(***A***) Estradiol levels after 4 weeks of E2 treatment in male and female NOD mice measured by ELISA. N=9 Placebo, 9 E2-treated mice. (***B***) Dot plots depicting expression of genes used to identify islet cell types in scRNAseq. The size of the dots indicates the percentage of cells that express the indicated gene. The color scale shows the change in normalized average gene expression within the different groups. N=3 biological replicates for scRNAseq. Statistical tests: Unpaired t-test: **A**. Data are represented as mean ± SEM.

